# Intrathecal antibodies cross-react with EBV BRRF2 and human antigens in multiple sclerosis

**DOI:** 10.64898/2026.08.14.744882

**Authors:** Poul M. Schulte-Frankenfeld, Tim Decker, Isabel Bünger, Sarah Bamberg, Manjusha Thakar, William R. Morgenlander, Patrick Schindler, Carolin Otto, Pia S. Sperber, Tanja Schmitz-Hübsch, Hans-Christian Kornau, Dietmar Schmitz, Sven Jarius, Erin E. Longbrake, Soumya Yandamuri, Kevin C. O’Connor, Carolin Schwake, Ilya Ayzenberg, Carlos A. Pardo, Friedemann Paul, Peter A. Calabresi, Klemens Ruprecht, H. Benjamin Larman, Jakob Kreye

## Abstract

Intrathecal antibody synthesis is a hallmark of multiple sclerosis (MS). Although some intrathecally synthesized antibodies in MS are known to target viral antigens, the spectrum of their antigenic specificities remains incompletely defined. We combined proteome-wide antibody profiling by Phage ImmunoPrecipitation Sequencing (PhIP-Seq) with cross-compartment analytics (MICAR) to study intrathecal antibody synthesis at peptide resolution in paired CSF and serum samples from individuals with MS (*n* = 40) and non-MS controls (*n* = 83).

While intrathecal antibody responses in MS were polyspecific and included reactivities to various viruses, we identified a subset of individuals with a convergent intrathecal antibody reactivity to a previously described motif within the Epstein-Barr virus (EBV) protein BRRF2 (BRRF2_408–415_). This motif-directed response showed co-reactivity with multiple CNS-expressed human antigens. In an independent cohort of *n* = 909 individuals with MS and *n* = 311 controls, including individuals with NMOSD and MOGAD, serum antibodies to BRRF2_408–415_ were detected in 8.36% of MS individuals and in 0.64% of non-MS controls, corresponding to an odds ratio for MS of 14.1 (95% CI: 4.4–86.04). Within MS individuals, BRRF2_408–415_ seropositivity was associated with increased intrathecal IgG synthesis. Cross-reactivity of antibodies to BRRF2_408–415_ with human targets, including TRIM71 and RTN2, was confirmed by competition ELISA and cell-based assays.

Together, these data define an intrathecal EBV BRRF2-linked antibody signature with human target cross-reactivity in a subset of MS individuals. This signature identifies a highly specific serological marker in MS and may support future stratification of the heterogeneous MS spectrum.

## Introduction

Multiple sclerosis (MS) is the most common chronic inflammatory disease of the central nervous system (CNS), affecting approximately 2.8 million individuals worldwide.^1^ MS is characterized by demyelinating lesions in various CNS regions.^2^ While its pathogenesis is incompletely understood,^3^ risk factors for MS include genetic susceptibility,^4–6^ environmental and lifestyle factors,^7–9^ and prior Epstein–Barr virus (EBV) infection.^10–12^ B-cells and antibodies play a central role in MS,^13–15^ as highlighted by the clinical efficacy of B-cell-depleting therapies^16,17^ and plasma exchange.^18,19^

A diagnostic hallmark of MS is intrathecal antibody synthesis (ITS). Here, antibodies prevalent in the cerebrospinal fluid (CSF) originate not only from passive diffusion across the blood–brain barrier, but also from local synthesis within the CNS.^20^ Intrathecal antibodies can be assessed by global measures, including the intrathecally synthesized antibody fraction and CSF-specific oligoclonal bands (OCBs),^21^ or by antigen-specific antibody indices, which are inherently restricted to selected predefined antigens.^22,23^ CSF-specific OCBs are detected in up to 95% of individuals with MS,^24,25^ but are rare in other demyelinating conditions such as neuromyelitis optica spectrum disorder (NMOSD) or MOG antibody-associated disease (MOGAD).^26,27^ Consistent with this clinical relevance, CSF-specific OCBs are an established component of MS diagnostic criteria,^28,29^ and are associated with a more severe disease course.^30,31^

Despite their diagnostic value and a growing body of evidence pointing towards a functional role of intrathecally synthesized antibodies in MS, their antigenic targets remain largely unidentified. Previous studies have described polyspecific intrathecal responses against common viruses such as measles, rubella, and varicella zoster — the so-called “MRZ reaction.” in individuals with MS.^32^ However, these ELISA-based studies are inherently limited, because they assess reactivity against predefined antigens. A complementary approach has involved recombinant expression of antibodies derived from CSF oligoclonal bands, revealing reactivity against ubiquitous self-proteins.^33^ However, this approach lacks scalability and generalizability for broader antigen discovery.

To overcome limitations in ITS characterization, we have recently developed a computational framework called Multiplexed Index Calculations of the Antibody Reactome (MICAR).^34^ This approach integrates paired CSF and serum antibody reactome profiles obtained via Phage ImmunoPrecipitation Sequencing (PhIP-Seq), enabling unselected proteome-wide quantification of intrathecal antibody synthesis.^35^ MICAR quantitatively distinguishes oligoclonal intrathecal antibody signals from polyclonal serum-derived background, resolving reactivity at the pathogen, protein, and peptide levels. Our initial study, which examined herpes simplex virus (HSV) responses in herpes simplex encephalitis (HSE) using samples from individuals with MS as controls, and did not investigate intrathecal antibody specificity in MS.^34^

Here, we applied MICAR in MS and control cohorts and identified a convergent intrathecal antibody response in a subset of individuals with MS. This shared reactivity is driven by a short motif in the EBV BRRF2 protein and cross-reacts with multiple human antigens expressed in the CNS.

## Results

### Intrathecal antibody responses in MS are polyviral with enhanced EBV reactivity

To characterize the antigenic specificity of intrathecally synthesized antibodies in MS, we analyzed paired CSF and serum reactivity profiles using multiple proteome-scale PhIP-Seq libraries (> 500,000 peptides in total; see *Material and methods*) from 40 individuals with MS and 83 non-MS controls with herpes simplex encephalitis (HSE) or anti-NMDA receptor encephalitis (NMDARE) (Supplementary Table 1). As expected, individuals with MS showed increased intrathecal antibody synthesis (ITS), reflected by oligoclonal band (OCB) frequency and the intrathecal antibody fraction (Supplementary Table 1). We first examined intrathecal antibody targets at the viral pathogen level. Individuals with MS and non-MS controls showed reactivity to a similar number of viral species in both serum and CSF (Fig. 1A and D). Consistent with prior work,^10^ EBV-specific reactivity was detected in all MS sera and CSF samples but was absent in 3/83 non-MS sera and 3/83 non-MS CSF samples (Supplementary Fig. 1B). Across > 200 viral species represented in the VirScan library, overall virus-specific binding intensities were largely comparable between MS and non-MS groups. The most notable difference was higher EBV reactivity in MS in both serum (Fig. 1B and C) and CSF samples (Fig. 1E and F).

**Figure 1:**
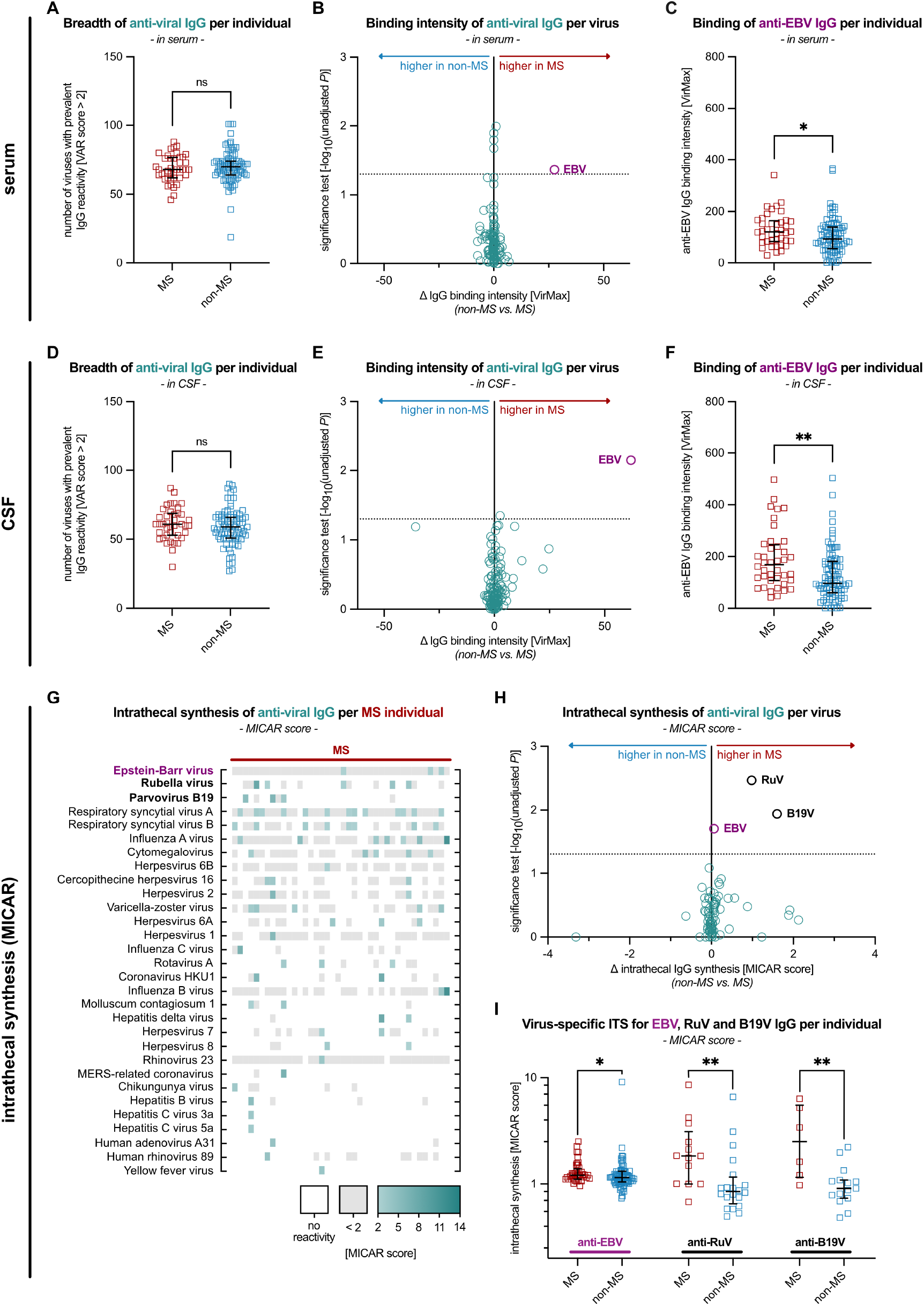
Viral antibody responses in serum, CSF and intrathecal synthesis. (**A**) Number of viruses with a detectable serum antibody response in each individual (VAR score > 2). Each square represents one individual (MS: *n* = 40, non-MS: *n* = 83). Group differences were assessed using a two-sided Mann–Whitney U test. Horizontal lines indicate the median; whiskers indicate the interquartile range. (**B**) Cohort-level differences in serum antibody binding intensity for each virus, measured as VirMax. Each circle represents one virus. Only viruses with detectable antibodies (VAR score > 2) in at least four MS samples (10% of the MS cohort; *n* = 129 viruses) are shown. The x-axis indicates the difference in median VirMax values between MS and non-MS; the y-axis shows unadjusted -log10(*P*) from a two-sided Mann–Whitney U test. (**C**) Individual-level EBV-specific serum antibody binding intensity, measured as VirMax, stratified by cohort. Each square represents one individual (MS: *n* = 40, non-MS: *n* = 83). Group differences were assessed using a two-sided Mann–Whitney U test. Horizontal lines indicate the median; whiskers indicate the interquartile range. (**D–F**) Corresponding analyses to (A-C) performed in CSF. (**G**) Individual-level heatmap of intrathecal virus-specific antibody synthesis in individuals with MS (*n* = 40), quantified by MICAR scores. Only viruses with ITS in at least one MS individual are shown. White indicates absence of detectable reactivity in CSF; grey indicates detectable but not intrathecally enriched reactivity; and green indicates intrathecally enriched reactivity, with darker shades corresponding to higher ITS magnitude. (**H**) Virus-level differences in ITS, measured by MICAR scores, between cohorts. Each circle represents one virus. Only viruses with ITS in at least one individual are shown (*n* = 77 viruses). The x-axis indicates the difference in median MICAR scores between cohorts; the y-axis shows unadjusted -log10(*P*) from a two-sided Mann–Whitney U test. (**I**) MICAR scores for EBV, rubella virus (RuV), and parvovirus B19 (B19V), stratified by cohort. Each square represents one individual with a computable virus-level MICAR score, defined by detectable CSF binding to ≥ 5 peptides of the respective virus (see *Material and methods*; EBV (MS: *n* = 40, non-MS: *n* = 75), RuV (MS: *n* = 13, non-MS: *n* = 20), B19V (MS: *n* = 6, non-MS: *n* = 15). Values are log-transformed. Group differences were assessed using a two-sided Mann–Whitney U test. Horizontal lines indicate the median; whiskers indicate the interquartile range.

We next quantified virus-specific intrathecal antibody synthesis. MICAR scores correlated with conventional ELISA-based antibody indices where available, supporting MICAR-based estimates of antiviral ITS (Supplementary Fig. 1C). Individuals with MS exhibited enhanced intrathecal reactivity to a median of two viral species per individual (IQR: 1.0–3.3), with up to seven distinct intrathecal viral responses (Fig. 1G), consistent with a polyviral ITS in MS.^36,37^ In contrast to HSE, where intrathecal responses were dominated by HSV and related viruses (Supplementary Fig. 1D), intrathecal viral responses in MS were heterogeneous (Fig. 1G). Significant differences in the magnitude of ITS between individuals with MS and controls were observed for three viruses (Fig. 1H): rubella virus (RuV; a component of the classic MRZ reaction),^32^ parvovirus B19 (B19V; recently reported as part of an expanded MRZ panel)^36^ and EBV. For rubella and parvovirus B19, the group-level differences were driven by subsets of MS samples with pronounced intrathecal reactivity. By contrast, EBV ITS showed no striking outliers; instead, EBV MICAR indices were only mildly but consistently increased at the group level in MS (median 1.2; IQR: 1.0–1.3) compared to non-MS controls (median 1.1; IQR: 0.9–1.2; Fig. 1I).

### Shared intrathecal antibody targets in an MS subgroup

Having established the virus-level context of intrathecal antibody responses, we next leveraged peptide-level MICAR scores across all PhIP-Seq libraries to identify shared intrathecal antibody targets in MS. We identified 27 peptides with significantly higher intrathecal enrichment in individuals with MS compared to non-MS controls (MS_ITS+_; Fig. 2A and Supplementary Table 4). To assess co-reactivity among these peptides, we computed a correlation network that revealed a central, highly connected cluster of 15 peptides (MS_ITS+ cluster_; Fig. 2B and Supplementary Table 4). UMAP embedding of intrathecal antibody reactivity from this cluster delineated a subgroup of six MS individuals with a distinct binding profile (Fig. 2C). These immunogenic cluster samples (MS_IC+_) showed frequent recognition of MS_ITS+ cluster_ peptides, whereas such reactivity was rare in the remaining MS and control samples (Fig. 2D and Supplementary Fig. 2A).

**Figure 2:**
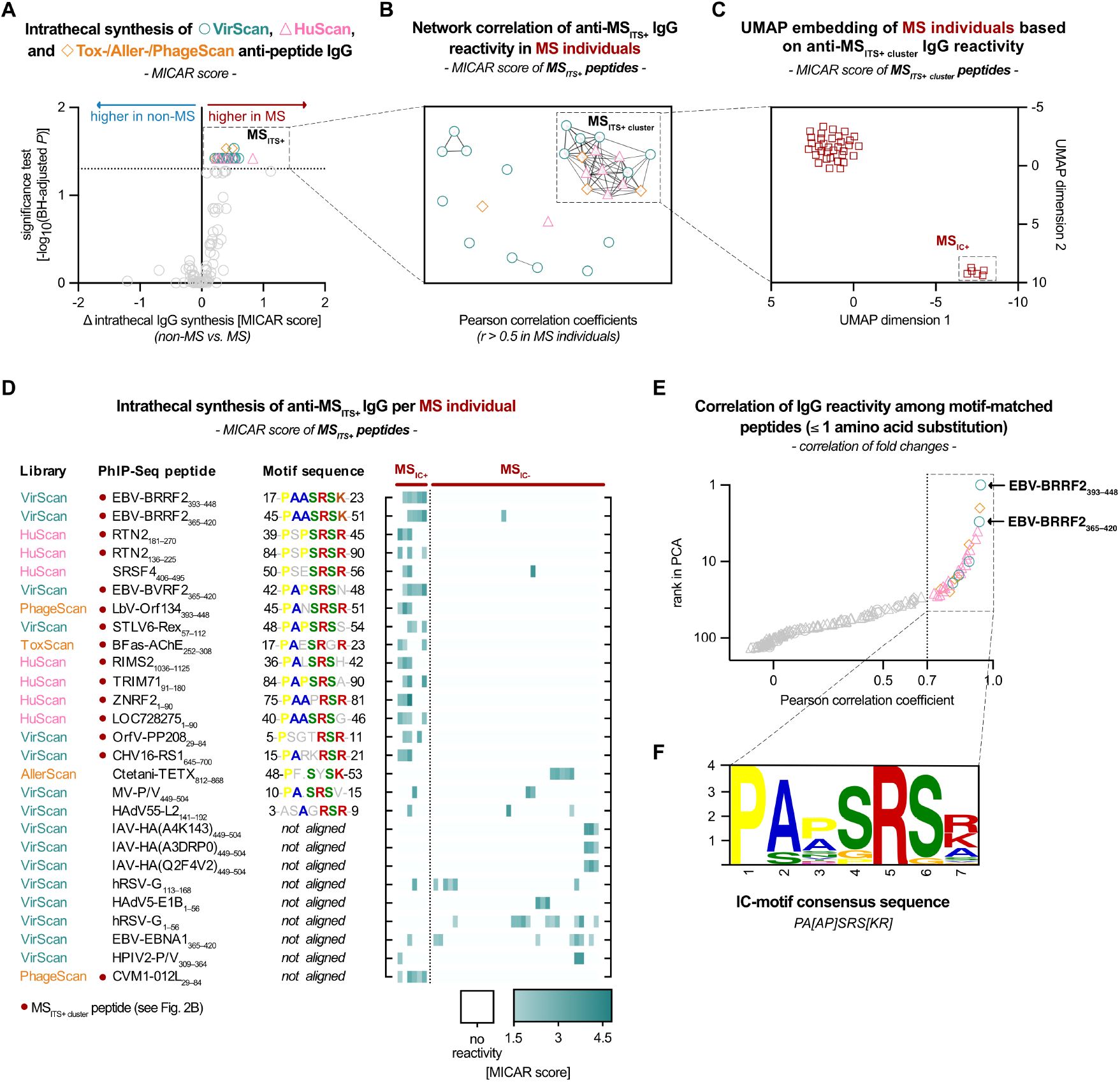
Peptide-level intrathecal antibody synthesis identifies an MS-associated immunogenic cluster. (**A**) Peptide-level differences in ITS, measured by MICAR scores, between MS and non-MS cohorts, shown as volcano plot. Each symbol represents one peptide, with only those shown that are detected in at least three MS individuals (*n* = 110 peptides). The x-axis indicates the difference in mean MICAR scores between MS and non-MS; the y-axis shows -log10(BH-adjusted *P*) from a two-sided Mann– Whitney U test with Benjamini–Hochberg (BH) correction. Colors and shapes denote PhIP-Seq libraries; non-significant peptides are shown in grey. Peptides with significantly greater CSF enrichment in the MS compared to the non-MS cohort (BH-adjusted *P* < 0.05) were classified as MS_ITS+_. (**B**) Correlation network of *n* = 27 MS_ITS+_ peptides based on Pearson correlation coefficients of MICAR scores across individuals with MS. Each symbol represents one peptide; edges represent correlations *r* > 0.5. The network was visualized using a Fruchterman–Reingold layout. Peptides within the primary network cluster were classified as MS_ITS+ cluster_ peptides. (**C**) UMAP embedding of MS individuals (*n* = 40) based on MICAR scores of *n* = 15 MS_ITS+ cluster_ peptides. Individuals assigned to the immunogenic cluster (IC) by *k*-means clustering in UMAP space were classified as MS_IC+_ (*n* = 6). (**D**) Individual-level heatmap of MICAR scores for MS_ITS+_ peptides in individuals with MS (*n* = 40), grouped by MS_IC_ status. White indicates absence of ITS; shades of green indicate magnitude of ITS. Motif sequences represent aligned central peptide regions. Colored residues indicate agreement with the IC-motif; grey residues indicate no agreement. (**E**) Correlation analysis of antibody binding among PhIP-Seq library peptides containing a motif with up to one amino acid mismatch to the IC-motif (*n* = 300 peptides) and binding in at least one sample (*n* = 137/300 peptides). Each symbol represents one peptide; colors and shapes are as in (A). The x-axis shows Pearson correlation coefficients for the five highest-ranking peptides across all samples; the y-axis indicates rank based on the first principal component. Peptides with correlations *r* ≥ 0.7 (*n* = 28) were reanalyzed for sequence alignment. (**F**) IC-motif derived from reanalyzed sequence alignment of peptides with correlating antibody binding (from E), shown as sequence logo (20% consensus). Letter height reflects relative amino acid frequency at each position.

Alignment of all MS_ITS+_ peptide sequences revealed a shared 7-amino-acid motif, consistent with the observed co-reactivity (Fig. 2D and Supplementary Fig. 2A). To further define the motif specificity, we evaluated all 300 motif-matched peptides across the PhIP-Seq libraries containing the motif with at most one amino acid substitution, of which 137 peptides bound in at least one sample, and correlated their reactivities with the five top-ranking MS_ITS+ cluster_ peptides (Fig. 2E, Supplementary Fig. 2B–C and Supplementary Table 5). Substitutions at positions 3, 4, or 7 frequently preserved correlation (*r* ≥ 0.7), whereas substitutions at other positions more often disrupted binding. We therefore used the subset of strongly correlating peptides (*r* ≥ 0.7) to refine the consensus, yielding the immunogenic cluster (IC) motif PA[AP]SRS[KR] (Fig. 2F). This motif closely resembles sequences independently reported as the target of antibodies in MS sera by prior studies.^38–40^

### EBV BRRF2 as the antigenic center of the motif-associated antibody response

Unlike earlier studies, which were limited to the human proteome, our analysis captured antibody reactivity to both human and viral antigens, enabling direct comparisons of antigenic contributors of the motif-associated response. In the motif-matched correlation analysis described above, two overlapping peptides from the EBV tegument protein BRRF2 — each containing an exact match to the IC consensus motif — were the top contributors to the principal component (Fig. 2E and Supplementary Table 5), whereas other previously proposed viral candidates ranked lower and showed weaker correlation with the MS_ITS+ cluster_ (Supplementary Fig. 2B–C and Supplementary Table 5).^40^ The two BRRF2 peptides were the only motif-matched sequences recognized by CSF antibodies from all MS_IC+_ individuals (Fig. 3A) and showed the strongest CSF binding and elevated MICAR indices in five of six MS_IC+_ individuals (Fig. 3A), nominating BRRF2 as the antigenically driving target of this antibody response.

**Figure 3:**
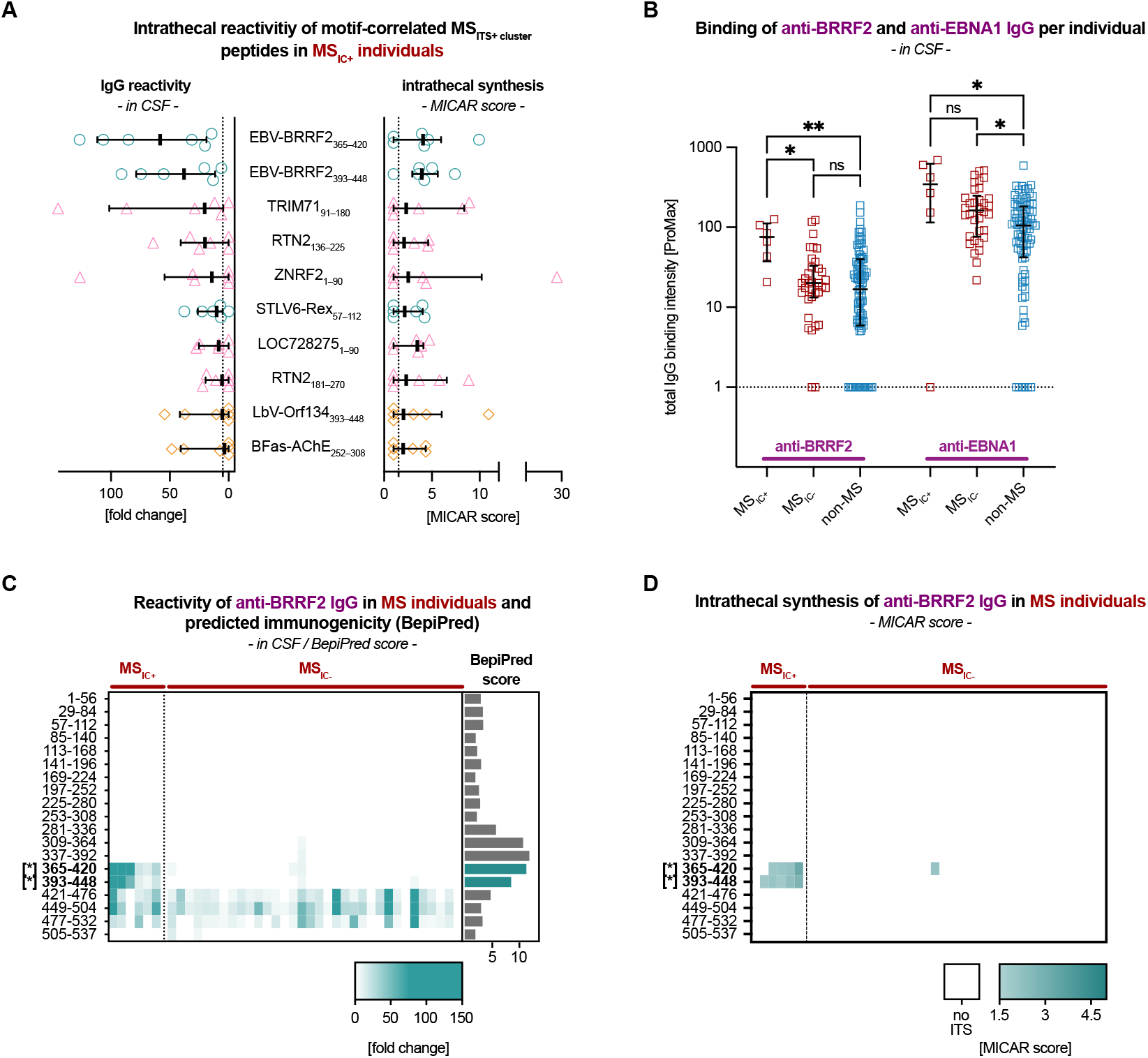
Characterization of intrathecal reactivity to motif-correlated MS_ITS+ cluster_ peptides. (**A**) Individual-level CSF reactivity and intrathecal synthesis for motif-correlated MS_ITS+ cluster_ peptides within the MS_IC+_ cohort (*n* = 6). Motif-correlated peptides were defined as those with correlation coefficients *r* ≥ 0.7 in the analysis shown in Fig. 2E. The left panel shows antibody binding intensity in CSF as fold change; the right panel shows the corresponding MICAR scores for each MS_IC+_ individual. (**B**) Individual-level CSF antibody binding intensity to BRRF2 and EBNA1, measured by ProMax, stratified by cohort. Each square represents one individual (MS_IC+_: *n* = 6, MS_IC-_: *n* = 34, non-MS: *n* = 83). Values > 1 indicate detectable reactivity. Group differences were assessed using a Kruskal–Wallis test (BRRF2: *P* < .05; EBNA1: *P* < .01) with Dunn’s post-hoc. Horizontal lines indicate the median; whiskers indicate the interquartile range. (**C**) Individual-level heatmap of CSF reactivity across the BRRF2 proteome. Rows correspond to separate BRRF2 tiles as represented in the PhIP-Seq library, positions within proteins indicated with asterisks indicating IC-motif-containing peptides. Columns represent individuals stratified by MS_IC_ status (MS_IC+_: *n* = 6, MS_IC-_: *n* = 34). White indicates absence of detectable reactivity; shades of green indicate magnitude of antibody binding intensity. BepiPred scores indicate predicted linear epitope immunogenicity (see Clifford JN et al. (2022) BepiPred-3.0: Improved B-cell epitope prediction using protein language models. *Protein Sci Publ Protein Soc*. doi:10.1002/pro.4497). (**D**) Individual-level heatmap of intrathecal synthesis across the BRRF2 proteome, quantified by MICAR scores. The layout of rows and columns corresponds to the heatmap in (C).

To further delineate the immunogenic basis of motif-containing antibody responses, we mapped reactivity across the full-length sequences of MS_ITS+ cluster_ proteins. For most motif-containing proteins, binding was largely restricted to the shared motif region (Supplementary Fig. 3A–G). In contrast, BRRF2 uniquely showed reactivity not only to the motif-containing region but also non-intrathecal reactivity to motif-independent adjacent sites, consistent with a prevalent, polyclonal response targeting this protein (Fig. 3C and Supplementary Fig. 3A–G). Sequence-based epitope prediction^41^ identified four consecutive BRRF2 peptides — including both IC-motif-containing peptides — as the most likely antibody epitope region (Fig. 3C, BepiPred). Despite this predicted immunogenicity, MS_IC-_ and non-MS samples showed BRRF2 reactivity restricted to non-motif regions (Fig. 3B and Supplementary Fig. 3A-G). By contrast, all MS_IC+_ individuals exhibited additional, intrathecally produced antibodies targeting the motif-containing region BRRF2_365–448_, resulting in increased overall BRRF2 reactivity without a comparable increase to other EBV proteins (Fig. 3B–D and Supplementary Fig. 3H). These data support an altered, intrathecal BRRF2-directed response as the antigenic center of the motif-associated (auto)reactivity.

### BRRF2_408–415_ antibodies are highly specific for MS and cross-react with human antigens

To further define antibody binding to the BRRF2-derived IC-motif and assess MS-specificity, we developed an ELISA-based assay amenable to clinical translation. Using sera with strong BRRF2_365–420_ reactivity in PhIP-Seq, we confirmed binding to a synthesized 56-mer BRRF2_365–420_ peptide that matched the PhIP-Seq library sequence (Supplementary Fig. 4A). Binding was markedly reduced when the IC-motif was scrambled or replaced with alanines, indicating that motif integrity is required for antibody recognition. Notably, the 7-mer motif alone (BRRF2_409–415_, PAASRSK) was not sufficient for binding, whereas extending the peptide by the preceding residue (S408) restored binding (BRRF2_408–415_;). ELISA reactivity to BRRF2_408–415_ correlated with PhIP-Seq signals in both serum and CSF (Supplementary Fig. 4B and C). When normalized to equal IgG concentrations, binding was stronger in CSF than in serum, consistent with intrathecal enrichment (Supplementary Fig. 4D).

To validate BRRF2_408–415_ antibodies in an independent cohort and estimate their prevalence, we analyzed a large multicenter cohort from the USA and Germany (Supplementary Table 2). Serum samples from individuals with MS (*n* = 909) and non-MS controls (*n* = 311), including individuals with NMOSD (*n* = 78) and MOGAD (*n* = 93), were tested for BRRF2_408–415_ antibodies using the established ELISA. As comparators, we also measured antibodies against a recently described EBNA1-motif (EBNA1_386–405_) and the related human cross-reactive antigen (GlialCAM_370–389_).^11^ BRRF2_408–415_ antibodies were detected in 8.36% of individuals with MS (*n* = 76/909) but were rare in non-MS controls (0.64%, *n* = 2/311), corresponding to a sensitivity of 8.36% and a specificity of 99.36% for MS in this cohort (Fig. 4A). This yielded an odds ratio (OR) of 14.1 (95% CI: 4.4–86.04; Fig. 4B), and the association remained robust after adjustment for age, sex and country of sampling in Firth penalized logistic regression (aOR: 9.48, 95% CI: 3.12–46.8); Supplementary Table 6). GlialCAM_370–389_ antibodies occurred at a similar frequency in MS (10.1%) but showed lower specificity (non-MS: 3.86%; specificity: 96.14%, OR: 2.81, 95% CI: 1.58–5.46). EBNA1_386–405_ antibodies were more common in the MS group (46.8%) than in the non-MS group (28.3%; specificity: 71.7%, OR: 2.23; 95% CI: 1.69–2.95). As expected, GlialCAM_370–389_ antibody reactivity was strongly associated with EBNA1_386–405_ reactivity (aOR: 19.72, 95% CI: 9.86–45.94, *p* < .0001), whereas BRRF2_408–415_ reactivity was largely independent of both (Supplementary Fig. 5A–C and Supplementary Table 7).

**Figure 4:**
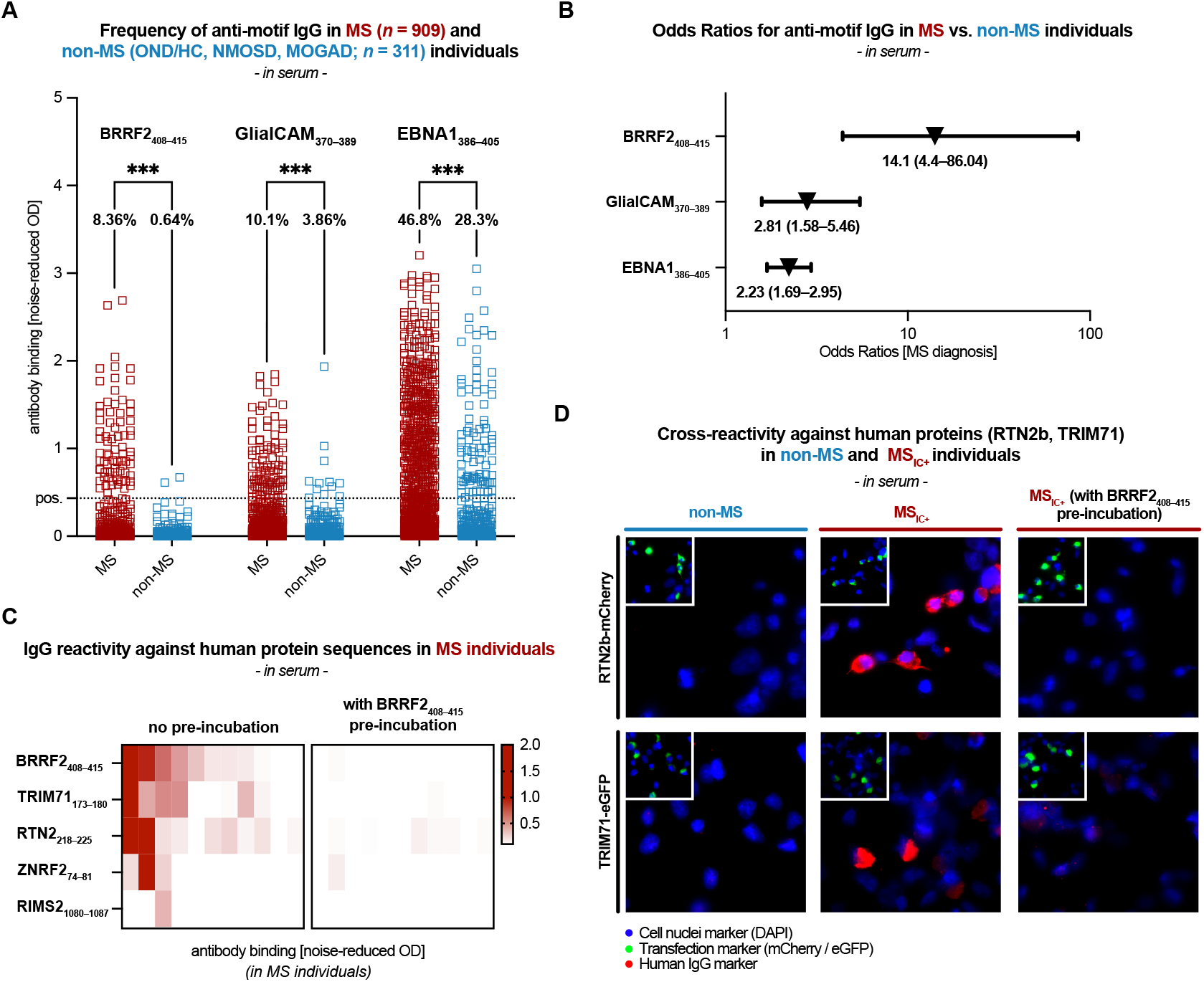
Experimental validation of the MS-associated IC-motif and its cross-reactivity with human antigens. (**A**) Individual-level ELISA-based IgG reactivity, measured as noise-reduced optical density (OD), against IC-motif BRRF2_408–415_, GlialCAM_370–389_, EBNA1_386–405_ in MS (*n* = 909) and non-MS (*n* = 311) cohorts. For validation of the positivity threshold (pos.) see *Material and methods*. Group differences were assessed using two-sided Mann–Whitney U tests. (**B**) Odds ratios for the association between IgG seropositivity to the peptides shown in (**A**) and MS, estimated using unadjusted logistic regression, with profile-likelihood 95% confidence intervals. (**C**) Individual-level heatmap of serum IgG reactivity against BRRF2_408–415_ and corresponding human IC-motif peptides derived from selected MS_ITS+ cluster_ peptides. Reactivity is shown before (left) and after (right) pre-incubation with BRRF2_408–415_. Columns represent individuals with MS (*n* = 11, including two from the discovery cohort). Red indicates increasing noise-reduced OD values. (**D**) Exemplary fluorescence microscopy images of cell-based assays using HEK cells overexpressing full-length human proteins RTN2b or TRIM71. The left column shows absence of serum antibody binding with a non-MS serum sample (representative of *n* = 5 non-MS samples tested); the middle column shows serum antibody binding with an MS_IC+_ serum sample (representative of *n* = 5 MS_IC+_ samples tested); and the right column shows blocked binding after pre-incubation of the same MS_IC+_ serum sample with soluble BRRF2_408–415_ peptide, as observed in all *n* = 5 MS_IC+_ samples tested.

Longitudinal serum samples were available for 108 individuals with MS, with a median interval of 1.1 years between the first and last sampling (IQR: 0.96–2.04, range: 0.14–9.96 years). Antibody levels were stable over time for BRRF2_408–415_ (ICC: 0.85, 95% CI: 0.79-0.9) and EBNA1_386–405_ (ICC: 0.89, 95% CI: 0.84–0.93), and moderately stable for GlialCAM_370-389_ (ICC: 0.59, 95% CI: 0.46–0.7). Binary serostatus at the first timepoint was positive in *n* = 14/108 individuals for BRRF2_408–415_, *n* = 9/108 for GlialCAM_370–389_, and *n* = 52/108 for EBNA1_386–405_, and remained highly concordant across timepoints with agreement rates of 95.4%, 94.4%, and 88.9%, respectively (Supplementary Fig. 5D-F).

To validate cross-reactivity, we tested MS_IC+_ sera against IC-motif peptides derived from selected motif-containing human proteins by ELISA, confirming heterogeneous binding patterns (Fig. 4C). Pre-incubation with the BRRF2_408–415_ peptide markedly reduced binding to these human motif peptides across all MS_IC+_ sera (Fig. 4C). For MS_IC+_ samples with highest ELISA binding (*n* = 5) and non-MS controls (*n* = 5), we next assessed whether these antibodies recognized full-length human proteins using cell-based assays with HEK293T cells overexpressing RTN2b and TRIM71, the strongest motif-associated targets from the PhIP-Seq and ELISA analyses. Antibody binding was observed in all MS_IC+_ sera but not in non-MS control sera and was strongly diminished by BRRF2_408–415_ peptide pre-incubation (Fig. 4D), confirming cross-reactivity in selected sera between BRRF2 IC-motif antibodies and motif-containing human proteins. Notably, several implicated human proteins show brain-enriched expression or genetic associations with neurological phenotypes (Supplementary Table 8), consistent with their potential functional relevance in MS.

### BRRF2_408–415_ antibodies associate with intrathecal IgG synthesis

After identifying a subset of MS individuals with IC-motif-binding intrathecal antibodies, we next assessed whether MS_IC+_ status was associated with clinical characteristics. Clinical data were available for the discovery cohort (*n* = 40) and for varying subsets of the validation cohort (*n* = 909 individuals with MS and *n* = 311 non-MS controls). MS_IC+_ and MS_IC−_ individuals did not differ significantly in age, sex, or basic CSF parameters (Table 1). However, MS_IC+_ individuals showed significantly increased intrathecal IgG synthesis, which may be partly explained by production of IC-motif antibodies. Intrathecal IgM synthesis was higher in MS_IC+_ individuals at a descriptive level, but this difference did not reach statistical significance (Supplementary Fig. 5G). Contrary to a previous report, exploratory analyses of age-adjusted serum neurofilament light chain (sNfL) z-scores as a marker of neuroaxonal damage did not differ between groups (Table 1 and Supplementary Fig. 5H).^40^

**Table 1.** Clinical and laboratory parameters in MS_IC+_ vs. MS_IC-_ from discovery and validation cohorts.

| Variables | MS<sub>IC+</sub> | MS<sub>IC-</sub> | non-MS | Sample size<sup>a</sup> | P-value<sup>b</sup><br>MS<sub>IC+</sub> vs.<br>MS<sub>IC-</sub> |
| --- | --- | --- | --- | --- | --- |
| **Number of individuals<sup>c</sup>** |  |  |  |  |  |
| from discovery cohort, n | 6 | 34 | n/a |  |  |
| from validation cohort, n | 76 | 833 | 309 |  |  |
| **Sample characteristics** |  |  |  |  |  |
| Age in years, median (IQR) | 47 (34–57) | 47 (37–56) | 48 (36–59) | [78, 810, 216] | .799 |
| Sex, %Female | 74.4% | 72.5% | 63.8% | [78, 810, 218] | .791 |
| Months since MS diagnosis, median (IQR) | 84 (47.5–192) | 108 (48–192) | n/a | [72, 766, n/a] | .214 |
| **Clinical characteristics** |  |  |  |  |  |
| EDSS, median (IQR) | 2 (1.5–3) | 3 (1.5–3.5) | 2.5 (1.5–4) | [68, 706, 69] | .115 |
| Serum NfL, pg/mL, median (IQR) | 11.3 (8–14.7) | 11 (8.7–15) | 10.7 (7.5–13.8) | [54, 646, 79] | .829 |
| Serum NfL age-adjusted z-score, median (IQR) | 0.6 (0–1.4) | 0.9 (0.2–1.5) | 0.3 (–0.5–0.9) | [54, 646, 79] | .161 |
| **CSF parameters** |  |  |  |  |  |
| CSF WBC, cells/μL, median (IQR) | 7 (2–15) | 6 (4–12) | 7.5 (2.3–13.8) | [19, 123, 30] | .859 |
| CSF total protein, mg/L, median (IQR) | 339 (270–408) | 352 (274–423) | 365 (334–442) | [6, 34, 22] | .806 |
| Age-adjusted QAlb, median (IQR) | 0.7 (0.4–0.9) | 0.7 (0.6–1) | 0.8 (0.7–1) | [19, 120, 17] | .176 |
| CSF-specific oligoclonal bands, % | 94.7% | 91.9% | 14.3% | [19, 123, 64] | >.999 |
| **Intrathecal Synthesis parameters** |  |  |  |  |  |
| Intrathecally synthesized IgG fraction, %, median (IQR) | 39.6 (0–48.9) | 4 (0–28) | 0 (0–0) | [19, 121, 26] | **.026\*** |
| Intrathecally synthesized IgA fraction, %, median (IQR) | 0 (0–0) | 0 (0–0) | 0 (0–0) | [19, 116, 19] | .908 |
| Intrathecally synthesized IgM fraction, %, median (IQR) | 0 (0–34.2) | 0 (0–2) | 0 (0–0) | [19, 119, 15] | .092 |

<sup>a</sup> Number of samples with available data for each respective variable. Format: [MS<sub>IC+</sub>, MS<sub>IC-</sub>, non-MS]
<sup>b</sup> Statistical test comparing MS<sub>IC+</sub> versus MS<sub>IC-</sub> with a Mann–Whitney U or Fisher's exact test as appropriate.
<sup>c</sup> MS<sub>IC+</sub> samples in the discovery cohort were identified as described in Fig. 2A–C. In the validation cohort, MS<sub>IC</sub> status is based on BRRF2<sub>408–415</sub> seropositivity as described in Fig. 4A. Two non-MS samples testing seropositive for BRRF2<sub>408–415</sub> were excluded from this analysis, see *Materials and methods*.
\* Statistical significance with $P < 0.05$ .
Abbreviations: CSF = cerebrospinal fluid; EDSS = Expanded Disability Status Scale; IgA/G/M = immunoglobulin A/G/M; IQR = interquartile range; MS = multiple sclerosis; MS<sub>IC</sub> = MS immunogenic cluster; NfL = neurofilament light chain; OCB = oligoclonal band; QAlb = albumin quotient; WBC = white blood cell count.

## Discussion

In this study we used unbiased, sequencing-based antibody reactome profiling with CSF-serum compartment analysis to map the proteomic targets of intrathecal antibodies in MS. We identified a subgroup of individuals with MS who showed convergent intrathecal reactivity to a short motif within the EBV protein BRRF2 and provided orthogonal evidence for cross-reactivity with multiple human antigens. In a large independent cohort, BRRF2_408–415_ seropositivity as measured by ELISA was rare in non-MS controls and thus highly specific for MS (OR: 14.1), supporting its relevance as MS-associated cross-reactive autoreactivity and as a candidate biomarker with potential utility in MS stratification.

These findings fit into a growing body of work linking EBV-directed antibody responses in MS to molecular mimicry with human antigens, including myelin basic protein (MBP), anoctamin-2 (ANO2), alpha-crystallin B chain (CRYAB) and glial cell adhesion molecule (GlialCAM).^11,42–44^ Most reported cross-reactive antibodies involve the latent EBV protein EBNA1 and have shown utility for identifying individuals with MS or at increased risk to develop MS.^45–47^ Cross-reactive antibodies against EBV proteins beyond EBNA1 — including LMP1, BFRF3 and BRRF2 — have also been proposed.^48,49^ By using a methodologically independent approach and focusing on convergent intrathecal antibody enrichment, we identified BRRF2_408–415_ motif antibodies, providing independent support for prior reports of this motif’s reactivity and relevance in MS.^38–40,50,51^ In contrast to prior motif-centered studies, our unbiased cross-compartment analysis directly localized the disease-associated intrathecal response to a BRRF2 motif epitope, nominating BRRF2_408–415_ as the antigenic centre of this response. This distinguishes BRRF2_408–415_-directed intrathecal reactivity from broader BRRF2 immunity and from EBNA1-associated motif responses. We then validated antibody cross-reactivity with self-proteins in competition experiments and cell-based assays, extending peptide-level observations to include binding of full-length human proteins in a cellular context. Although our data nominate BRRF2 as the antigenic centre of the motif-associated response, they do not establish whether BRRF2 initiated this response or instead represents a highly immunogenic member of a broader shared motif reactivity. Longitudinal antibody studies will be required to distinguish these possibilities.

Among the human cross-reactive targets, BRRF2_408–415_ antibodies showed heterogeneous binding to four CNS-expressed human proteins, including RTN2, TRIM71, ZNRF2 and RIMS2. These targets are linked to neuronal structure, RNA regulation, ubiquitin-dependent signalling, microglial biology or synaptic vesicle function, and several have genetic associations with neurological phenotypes.^52–57^ Although these observations support biological plausibility, the targets are predominantly intracellular, and whether antibodies access their native epitopes in vivo remains unresolved. Nonetheless, intracellular antibody uptake has been reported in other autoimmune conditions, including myositis, with demonstration of biological effects consistent with target interference.^58–60^ In MS, future studies using patient-derived monoclonal BRRF2_408–415_ antibodies can clarify their ability to access native CNS targets in situ, exert pathogenic or modulatory effects, interfere with synaptic transmission, post-transcriptional regulation, or alter cellular homeostasis.^81^ If pathogenicity is established, IC-motif antibodies may represent a therapeutic target for selective antibody depletion or B-cell eradication,^93,94^ and should be further considered in ongoing EBV vaccine development efforts.^61,62^

Beyond the observed BRRF2_408–415_-centered reactivity, we observed polyviral intrathecal antibody responses in MS, including elevated responses to rubella virus and parvovirus B19 in subsets of individuals. The latter may motivate explorations to expand the MRZ reaction, an established marker of polyspecific intrathecal virus antibody synthesis in MS, to enhance diagnostic sensitivity.^36^ Despite this pronounced polyviral intrathecal response and the strong epidemiological link between MS and EBV immunity, we did not observe a dominant, globally elevated intrathecal EBV response. The universal EBV seroprevalence and the only mild intrathecal antibody synthesis to EBV are consistent with prior observations and have been referred to as the “EBV paradox”.^63–66^ One possible explanation is that intrathecal synthesis in MS reflects persistence of B-cells that were nonspecifically recruited to the CNS, for example during primary EBV infection,^37,63,67^ with EBV subsequently promoting their long-term maintenance via modulation of host immune signaling in infected cells.^68^ This could favor intrathecal accumulation of diverse, polyviral B-cell populations, suggesting that EBV may contribute to MS not through the overall magnitude of the intrathecal EBV-directed antibodies, but through shaping the persistence, long-term survival, or autoreactive potential of selected B-cell clones, perhaps via cellular infection by EBV.^68–70^ In this framework, BRRF2_408–415_-reactive clones may represent one such example, although their actual temporal dynamics and affinity maturation trajectory after EBV infection remain to be determined experimentally.

Meanwhile, the factors initiating autoreactivity in MS remain incompletely understood. Prior work implicates altered immune control of EBV^71,72^ and host immune genetics — particularly HLA variation — associated with impaired viral clearance and reduced elimination of autoreactive GlialCAM_370–389_ B-cells.^73,74^ Carrying HLA-DRB1*15:01 amplifies the MS risk associated with EBNA1-motif seropositivity, supporting a joint contribution of host genetic variation and EBV infection to disease pathogenesis.^45^ Consistent with this concept, EBV infection can reprogram antigen presentation in HLA-DRB1*15:01-positive B-cells toward myelin basic protein (MBP)-derived self-peptides.^75^ Taken together, these observations warrant further investigation into whether BRRF2_408–415_-seropositivity is influenced by HLA background.

Differential EBV strain-related pathogenicity may represent an additional risk factor. Although EBV genes show substantial polymorphism, only a single BRRF2 missense variant has been reported to occur more frequently in MS than in control isolates.^76^ Of interest, this substitution (S412Y) is located within the BRRF2_408–415_ motif. Additional synonymous or non-coding variants within EBV genes are more frequently found in individuals with MS, including one spanning BRRF2- and EBNA1-related loci (g.94289G>T).^77^ Such sequence variation could plausibly alter epitope conformation, latency regulation, or immunogenicity, thereby shaping the selection of motif-reactive antibodies in genetically susceptible hosts, consistent with emerging evidence that HLA-associated antibody specificities are conditioned by intrinsic antigen properties.^78–81^ Thus, EBV strain analyses may be informative and could be revisited in HLA- and motif-stratified cohorts.

A key strength of this work is the translation of this intrathecally defined signature into a scalable serum ELISA and its validation in a large multicenter cohort. BRRF2_408–415_ seropositivity was detected in individuals sampled at variable times after symptom onset and across differing levels of disease activity,^50^ suggesting that IC-motif reactivity may represent a relatively stable serological state with higher disease specificity than any of the single comparator antibody biomarkers measured here and in prior work.^45,46^ At the same time, the restricted prevalence raises the possibility that BRRF2_408–415_ antibodies mark a biologically distinct MS endophenotype,^82,83^ or, alternatively, even a distinct inflammatory demyelinating patient group, analogous to how aquaporin-4 and MOG antibodies led to recognition of NMOSD and MOGAD as distinct disease entities.^84^ Notably, our analysis of clinical characteristics showed significantly increased intrathecal IgG synthesis, but no difference in sNfL levels in BRRF2_408–415_ seropositive individuals, contrasting findings from prior work.^40^ Hence, detailed phenotypic exploration in ideally longitudinal cohorts will be needed to test whether BRRF2_408–415_ seropositivity — and potentially seropositivity to EBNA1-related motifs — likewise associates with phenotype, disease progression, or treatment response.

The findings of the present study are limited by the restricted repertoire of linear epitopes represented in the PhIP-Seq library, which does not capture conformational or post-translationally modified epitopes. The selection of individuals with HSE and NMDARE as inflammatory comparator group in the relatively small, paired CSF–serum discovery cohort may have additionally influenced virus-level comparison due to the specific immunological characteristics of HSE/NMDARE patients. To compensate, the serum validation cohort primarily assessed specificity against other demyelinating and non-demyelinating conditions. Given that BRRF2_408–415_ antibodies are synthesized intrathecally, the restriction of the validation cohort to serum samples might underestimate the prevalence of this antibody response in the broader MS population. Treatment status at sampling was not systematically modelled and may have influenced serum antibody levels or clinical association analyses. Additional control conditions and complementary approaches using technologies such as molecular indexing of proteins by self-assembly (MIPSA) or rapid extracellular antigen profiling (REAP), as well as profiling of additional immunoglobulin classes, may therefore provide further insight.^85,86^

Clinically, despite the relatively low seroprevalence of ∼8% in the MS population, the much lower frequency of BRRF2_408–415_ antibodies in disease-relevant controls supports their potential utility as a high-specificity blood-based biomarker. Rather than serving as a broadly sensitive diagnostic marker, BRRF2_408–415_ seropositivity may help to stratify biologically defined MS subgroups and complement CSF-based measures of ITS used in MS diagnosis, most prominently OCBs and — in recent updates to the McDonald diagnostic criteria — CSF kappa free light chains (kFLC).^87^ Because BRRF2_408–415_ is detectable in serum and reflects a distinct EBV-directed response, it may provide a serum-based readout linked to ITS in a subset of individuals with MS. Finally, BRRF2_408–415_ could be incorporated into composite EBV immune signatures with low sensitivity but high specificity, which may improve risk stratification in preclinical or early MS cohorts, for example following infectious mononucleosis.^88^ Given that EBV infection, biomarker evidence of neuroaxonal injury and signature antibody presence can precede the clinical diagnosis of MS by ∼5–10 years,^10,40^ such markers may in future enable identification of high-risk individuals before symptom onset and thereby shift “hit hard and early” strategies toward earlier, risk-adapted intervention.^89,90^

## Materials and methods

### Study approval

This study was approved by the Ethics Committee of Charité – Universitätsmedizin Berlin, corporate member of Freie Universität Berlin and Humboldt–Universität zu Berlin, Germany (ethical vote number EA4/046/23). Biospecimen used in this study were sampled in multiple studies. Written informed consent was obtained from participants at all recruiting study centers.

### Patient cohorts & biospecimens

Discovery cohort (Fig. 1–3): Paired CSF and serum samples for the discovery cohort were collected for a study on anti-N-methyl-D-aspartate receptor (NMDAR) encephalitis (NMDARE) following herpes simplex encephalitis (HSE), in which individuals with MS served as a control cohort.^34^ Here, we reanalyzed the corresponding dataset focusing on convergent responses in the MS cohort. Samples of *n* = 40 individuals with MS diagnosed according to McDonald 2017 criteria were obtained from the Central Biobank of Charité – Universitätsmedizin Berlin (ethical vote EA4/018/17).^28^ As non-MS controls, we used paired CSF and serum samples from individuals with NMDARE or HSE (total *n* = 83; thereof *n* = 5 with HSE only (H_only_), *n* = 65 with NMDARE only (N_only_), *n* = 13 with HSE and secondary NMDARE (H^+^N^+^)), recruited across 16 centers participating in the German Network for Research on Autoimmune Encephalitis (GENERATE; ethical vote EA1/258/18), as well as from Johns Hopkins School of Medicine, Department of Neurology. Sample characteristics of the discovery cohort are available in Supplementary Table 1.

Validation cohort (Fig. 4): For validation, additional serum samples from individuals with MS (*n* = 909, including 108 individuals with a longitudinal follow-up sample) and non-MS controls (*n* = 311 individuals) were collected across two centers in the US (Johns Hopkins School of Medicine and Yale University School of Medicine) and two centers in Germany (Charité – Universitätsmedizin Berlin, including the BERLImmun study cohort,^91^ and Ruhr University Bochum; ethical votes EA1/182/10, EA1/163/12 and EA1/362/20). For individuals with longitudinal sampling, cross-sectional analyses were based on the earliest available sample. The non-MS controls were individuals with NMOSD (*n* = 78, including *n* = 6 individuals tested negative for AQP4-antibodies),^43,44^ MOGAD (*n* = 93),^92^ and non-demyelinating controls (OND/HC; *n* = 140), comprising other neurological disease controls (OND), including patients with idiopathic headaches, migraines, functional neurological disorders, idiopathic intracranial hypertension, normal pressure hydrocephalus, and healthy controls (HC). See Supplementary Table 2 for demographic characteristics of the validation cohort. Serum samples were processed and stored frozen at -80°C according to site-standard protocols, and were shipped and analyzed under pseudonymized identifiers according to study specific protocols. The laboratory personnel remained blinded to individual characteristics until completion of the PhIP-Seq experiments and primary analyses.

### PhIP-Seq experiments & MICAR algorithm

For IgG profiling, PhIP-Seq libraries displaying tiled proteomes of humans (HuScan; 274,207 90-mer peptides),^35,93^ human viruses (VirScan; 110,215 56-mer peptides), allergens (AllerScan; 19,331 56-mer peptides),^94^ environmental toxins and virulence factors (ToxScan; 95,601 56-mer peptides),^95^ and bacteriophages (PhageScan; 100,275 56-mer peptides)^96^ were used in a combined assay according to established protocols,^97^ as described previously for this dataset.^34^ In short, PhIP-Seq data were analyzed as fold changes relative to “beads-only” mock immunoprecipitation controls. The following classifiers for reactive “hits” were applied: Peptides with ≥ 15 counts, edgeR -log_10_(*P*) ≥ 3 compared to beads-only controls and a fold change of ≥ 5 compared to beads-only controls. Unless otherwise specified, downstream analyses were performed on fold change values of peptides scored as hits, with non-reactive peptides assigned a value of 1. For analyses of antibody responses on the level of viruses and proteins, aggregated scores were used. Viral Aggregate Reactivity (VAR) scores were computed by comparing average fold-change binding of peptides associated with each virus versus distributions of randomly selected peptides, and used to qualitatively compare discrete virus reactivity between individuals. Following previously established cut-offs, VAR scores ≥ 2 were interpreted as positive reactivity.^98–100^ VirMax scores were defined as the mean reactivity of the five most reactive peptides from five independent proteins per virus and used to quantitatively compare virus-level reactivity. ProMax scores were defined as the highest measured reactivity across all peptides mapped to a given protein, and used to quantitatively compare protein-level reactivity.

Intrathecal antibody synthesis was assessed using the Multiplexed Index Calculations of the Antibody Reactome (MICAR) algorithm, which quantifies relative antibody enrichment in CSF compared with paired serum based on normalized fold changes of IgG binding to PhIP-Seq peptides. Virus-level MICAR scores were computed from peptide-level MICAR scores and have been previously defined as the mean of peptide-level MICAR antibody indices for viruses with ≥ 5 peptide hits detected in the CSF. Virus MICAR scores > 1 indicate relatively greater CSF binding, and scores ≥ 2 were classified as intrathecal synthesis. Because MICAR scores indicate intrathecal enrichment rather than absolute CSF binding intensity, strong CSF reactivity can yield MICAR scores near 1 when comparable reactivity is also present in serum. A detailed description of the MICAR metrics, including validation procedures and cohort-specific quality control measures, has been published previously.^34^

### Motif discovery and peptide sequence comparison

Co-reactivity among peptides with more prevalent intrathecal synthesis in MS compared to non-MS (MS_ITS+_ peptides) was assessed using Pearson correlation across MS samples. Correlations > 0.5 were used to construct a weighted, undirected network using R {igraph} and {ggraph} (Fig. 2B).^101,102^ MICAR scores of MS_ITS+ cluster_ peptides were embedded using UMAP (R package {uwot}; *n_neighbors* = 5, *min_dist* = 0.2, *metric* = cosine, *seed* = 289). K-means clustering was subsequently applied to the two-dimensional embedding (*k* = 2) (Fig. 2C).^103^

The convergent intrathecal antibody response to the immunogenic cluster (IC) motif was identified using the GLAM2 gapped motif algorithm.^104,105^ An initial consensus motif (P[AS][PAE]SRS[RK]), retaining amino acids present in at least 20% of aligned sequences at each position, was identified from MS_ITS+_ peptides and mapped to 300 peptides across PhIP-Seq libraries containing the motif with at most one amino acid substitution. Relative binding scores of 137/300 peptides which bound in at least one sample (counts ≥ 15) across all CSF and serum samples from MS and non-MS cohorts were submitted to a Principal component analysis (PCA). Motif correlation scores, defined as the mean correlation of each peptide to the five highest ranking peptides in the first principal component of the PCA (see Supplementary Table 5), were calculated. Peptides with motif correlation scores *r* ≥ 0.7 were re-submitted to GLAM2, yielding a refined 20% consensus motif (P[AS][PA]SRS[RK]). Predicted immunogenicity of PhIP-Seq peptides (Fig. 3C) was computed using BepiPred-3.0 scores by summing predictions across all residues within each peptide.^41^

### Experimental validation by ELISA

For orthogonal validation, peptides representing IC-motif variants derived from BRRF2, EBNA1 and motif-containing human proteins were synthesized at ≥ 85% purity by the Charité Institute of Biochemistry using standard solid-phase peptide synthesis. All peptides were synthesized with a free, non-amidated C-terminus and carried a C-terminal His-tag followed by Strep-tag II (“His-Strep-tag”). A full list of sequences is provided in Supplementary Table 3.

Peptide ELISAs were performed on high-binding 96-well plates (Greiner Bio-One, #655101) coated overnight at 4°C with 2 µg/ml of the respective peptide in PBS. Plates were subsequently washed (PBS containing 0.05% Tween 20) and blocked with blocking buffer (PBS containing 0.1% BSA and 0.05% Tween 20). Serum (1:200) and CSF (1:20) samples were diluted in blocking buffer, then incubated for 2 hours at room temperature, then washed, and subsequently incubated with anti-human IgG MT78-ALP (1:1000; Mabtech #3850-9A) for one hour. Plates were washed and developed using dPNPP substrate (Thermo Fisher, #37620) and optical density (OD) was measured at 405 nm after 30 minutes. For inhibition experiments, sera were pre-incubated overnight at 4°C with the BRRF2_408–415_ peptide at 20 µg/ml in blocking buffer prior to the ELISA. Total IgG levels were assessed in parallel using anti-human IgG (MT145) coated wells (serum dilution 1:100,000; Mabtech #3850-1-250) to control for non-specific IgG loss. All ELISA measurements were performed in triplicate, summarized as the median and noise-corrected by subtracting sample-specific binding signals to His-Strep-tag control peptides (noise-reduced OD).

To define the seropositivity cut-off for the peptide ELISA experiments (Fig. 4A and B), sixteen EBV-seronegative non-MS serum samples (EBV VAR score < 2) from the discovery and validation cohorts (NMDARE: *n* = 6, MOGAD: *n* = 3, NMOSD: *n* = 1, OND/HC: *n* = 6) were assessed. Mean noise-reduced ODs of these samples to the BRRF2_408–415_, GlialCAM_370–389_ and EBNA1_386–405_ peptides were near zero (range: -0.007–0.007) with standard deviations ≤ 0.05 (range: 0.03–0.05) for all three peptides. Because background reactivity was near zero and comparable across peptides, seropositivity was defined using a uniform threshold of 0.433, corresponding to the pooled mean noise-reduced OD plus 10 pooled standard deviations (*mean_pooled_* = 0.0002, *SD_pooled_* = 0.0433).

### Cell-based assays

Cell-based assays were performed using HEK293T cells transiently overexpressing RTN2b (Addgene #186600, mCherry) or TRIM71 (VectorBuilder #VB900022-0960sgb, eGFP). Cells were fixed 48 hours post-transfection with methanol at -20°C for 4 minutes, washed, then blocked and permeabilized in blocking solution (PBS containing 5% normal goat serum, 2% bovine serum albumin and 0.1% Triton X-100).^106^ Cells were then incubated overnight at 4°C with sera (1:200 in blocking solution). IgG binding was detected using an anti-human IgG antibody (1:1000 in blocking solution) labelled with Alexa Fluor 488 (Dianova, #109-545-003) for RTN2b and Alexa Fluor 647 (BioLegend, #405322) for TRIM71, respectively. In Fig. 4D, for better comparability, both transfection markers (eGFP and mCherry) are visualized in green and human IgG binding (Alexa FluorFlour 488 and Alexa Fluor 647) is visualized in red. Nuclei were stained using 4′,6-diamidino-2-phenylindole (DAPI) and, in Fig. 4D, visualized in blue. For the BRRF2_408–415_-incubated condition, sera were pre-incubated at 4°C overnight with BRRF2_408–415_ peptide prior to staining of cells. Images were acquired using fluorescence microscopy (Leica SPE) with identical recording conditions for all images.

### Clinical data collection

For comparative analyses, MS_IC+_ status was defined primarily by MICAR positivity to the IC-motif in paired CSF–serum PhIP-Seq data from the discovery cohort. In the serum-only validation cohort, BRRF2_408–415_ seropositivity by ELISA was used as a surrogate marker of the IC-motif-associated antibody response. Routine diagnostic parameters, Expanded Disability Status Scale (EDSS) scores and clinical symptom status were obtained from medical records and study databases.^107,108^ OCB patterns type 2 and 3 were classified as CSF-specific according to consensus criteria.^109^ Age-adjusted albumin quotients (QAlb) were calculated by dividing the measured QAlb by the age-adjusted upper reference limits, with values >1 indicating blood-CSF barrier dysfunction.^22^ Serum neurofilament light chain (sNfL) concentrations were measured using the Atellica IM Neurofilament Light Chain assay. Age-adjusted z-scores were calculated using the Serum Neurofilament Light Measurement App, with the Body Mass Index set to 25kg/m^2^ for all individuals.^110^ The underlying reference cohort was measured with the Quanterix Simoa NF-Light V2 Advantage (PLUS) assay; however, given the strong association between Atellica and Simoa measurements (*R²* ≥ 0.95) and our restriction to internal subgroup comparisons within a uniformly Atellica-measured cohort, the app-based z-scores were considered appropriate in the absence of an assay-specific reference dataset.^111^

### Statistical analysis

Statistical analyses and figure generation were performed using RStudio (v2024.04.2) and GraphPad Prism (v10.6.1).^112^ Unpaired data were analyzed using unpaired *t*-tests or Mann–Whitney U tests with Benjamini-Hochburg (BH) correction where appropriate, paired data using paired *t*-tests or Wilcoxon signed-rank tests, and comparisons involving more than two groups using Kruskal–Wallis tests with Dunn’s correction, as appropriate. Categorical data were analyzed using Fisher’s exact tests. Correlations were assessed using simple linear regression. Odds ratios (OR) were estimated using unadjusted logistic regression with profile-likelihood 95% confidence intervals (95% CI) and Firth penalized logistic regression with profile penalized-likelihood 95% CI. Adjusted Firth penalized logistic regression odds ratio (aOR) models included age at sampling, sex and country of sampling, and models of inter-antibody association were additionally adjusted for disease group. All adjusted analyses were restricted to complete cases across all model variables. Intraclass correlation coefficients (ICC) were calculated using a two-way random-effects model for absolute agreement on single measurements. All *P*-values were two-tailed and considered significant at * *P* < 0.05, ** *P* < 0.01, *** *P* < 0.001 and **** *P* < 0.0001.

## Abbreviations

EBV: Epstein-Barr virus
EDSS: Expanded Disability Status Scale
ELISA: enzyme-linked immunosorbent assay
HC: healthy controls
HSE: herpes simplex encephalitis
HSV: herpes simplex virus
IC: immunogenic cluster
ITS: intrathecal synthesis
MICAR: Multiplexed Index Calculations of the Antibody Reactome
MOGAD: myelin oligodendrocyte glycoprotein antibody-associated disease
MRZ: measles-rubella-zoster
MS: multiple sclerosis
NfL: neurofilament light chain
NMDARE: anti-NMDA receptor encephalitis
NMOSD: neuromyelitis optica spectrum disorder
OCB: oligoclonal bands
OD: optical density
aOR: adjusted Odds Ratios
PCA: principal component analysis
PhIP-Seq: Phage ImmunoPrecipitation Sequencing
UMAP: Uniform Manifold Approximation and Projection

## Data availability

The authors confirm that all data supporting the key findings of the analyses presented in this study are available within the article and its supplementary data. Derived data required to reanalyze the reported findings of this study are available from the corresponding authors upon request and subject to applicable ethical and data protection requirements.

## Acknowledgements

The authors are grateful to all patients, their caregivers and to all clinical investigators contributing to the present study. The authors further thank Stephen Elledge, PhD, for providing the VirScan and human proteome PhIP-Seq libraries.

## Supplementary material

Supplementary material is available online.

