## Supplementary for "Intrathecal antibodies cross-react with EBV BRRF2 and human antigens in multiple sclerosis"

**Preprint: Supplementary Material for**

Department of Pediatric Neurology, Charité – Universitätsmedizin Berlin, corporate member of Freie Universität Berlin, Humboldt-Universität zu Berlin, and Berlin Institute of Health, Augustenburger Platz 1, 13353 Berlin, Germany.

and H. Benjamin Larman, Ph.D.

Johns Hopkins School of Medicine, Miller Research Building, Room 607, 733 North Broadway, Baltimore, MD 21205-1832

**Author affiliations:**

1 Charité – Universitätsmedizin Berlin, corporate member of Freie Universität Berlin and Humboldt Universität zu Berlin, Department of Pediatric Neurology, Berlin, Germany

2 Berlin Institute of Health at Charité – Universitätsmedizin Berlin, Germany

3 Department of Surgery – Minimally Invasive and Visceral Surgery, Vivantes Klinikum Neukölln, Berlin, Germany

4 Institute for Cell Engineering, Division of Immunology, Department of Pathology, Johns Hopkins University School of Medicine, Baltimore, MD, USA

5 Charité – Universitätsmedizin Berlin, corporate member of Freie Universität Berlin and Humboldt-Universität zu Berlin, Department of Neurology and Experimental Neurology, Berlin, Germany

6 German Center for Neurodegenerative Diseases (DZNE) Berlin, Berlin, Germany

7 Experimental and Clinical Research Center, a cooperation between Max Delbrück Center (MDC) for Molecular Medicine in the Helmholtz Association, Charité – Universitätsmedizin Berlin, Berlin, Germany

8 Charité – Universitätsmedizin Berlin, corporate member of Freie Universität Berlin and Humboldt-Universität zu Berlin, Institute of Medical Immunology (IMI), Berlin, Germany

9 Charité – Universitätsmedizin Berlin, corporate member of Freie Universität Berlin and Humboldt-Universität zu Berlin, Neuroscience Research Center (NWFZ), Berlin, Germany

10 Division of Neuroimmunology, Department of Neurology, University of Heidelberg, Heidelberg, Germany

11 Department of Neurology, Yale University School of Medicine, New Haven, CT, USA

12 Department of Immunobiology, Yale University School of Medicine, New Haven, CT, USA

13 Ruhr University Bochum, Department of Neurology, St. Josef Hospital, Bochum, Germany

14 Department of Neurology, Johns Hopkins University School of Medicine, Baltimore, MD, USA

### SUPPLEMENTARY FIGURES

##### Supplementary Figure 1: Viral antibody responses in serum, CSF and intrathecal synthesis

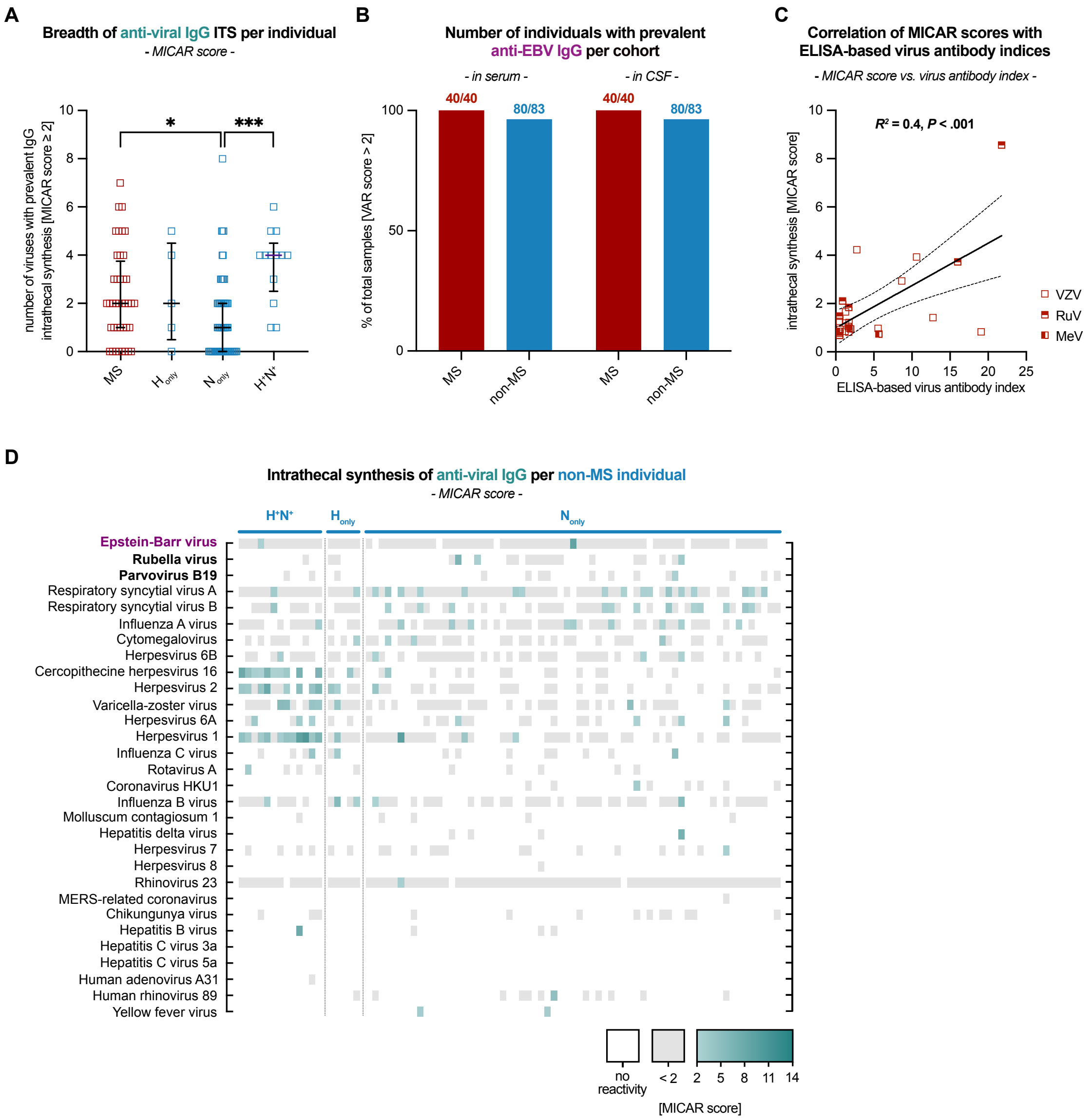


**(A)** Number of viruses with detectable intrathecal antibody enrichment in each individual (MICAR score > 2). Each square represents one individual in the MS cohort ($n$ = 40) and non-MS subgroups ($H_{only}$: $n$ = 5, $N_{only}$: $n$ = 65, $H^+N^+$: $n$ = 13). Group differences were assessed using a Kruskal–Wallis test ($P$ < .0001) with Dunn's post-hoc, only significant post-hoc tests are indicated by asterisks; horizontal lines indicate the median, whiskers indicate the interquartile range.

(**B**) Number of individuals with a detectable response against Epstein-Barr Virus (EBV) peptides (VAR score > 2) per cohort in serum and CSF.

(**C**) Individual-level intrathecal antibody enrichment quantification measured using MICAR scores, correlated with corresponding intrathecal antibody enrichment measures from routine clinical ELISA-based antibody indices for MRZ viruses: measles virus (MeV), rubella virus (RuV) and varicella-zoster virus (VZV), in individuals with MS (MeV: $n$ = 3, RuV: $n$ = 7, VZV: $n$ = 15). Each square represents one individual's measurement for a given virus, some individuals had data available for multiple viruses. The x-axis shows the ELISA-based clinical antibody index; the y-axis shows the MICAR score. MICAR scores and ELISA-based antibody indices are positively associated in a linear regression model ($R^2$ = 0.4, $P < .001$). Dashed lines indicate 95% confidence intervals.

(**D**) Extension of Fig. 1G. Individual-level heatmap of intrathecal virus-specific antibody synthesis in non-MS individuals (total: $n$ = 83; thereof $H^+N^+$: $n$ = 13, $H_{only}$: $n$ = 5, $N_{only}$: $n$ = 65), quantified by MICAR scores. Only viruses with ITS in at least one MS individual are shown. White indicates absence of detectable CSF reactivity; grey indicates detectable but not intrathecally enriched reactivity; and green indicates intrathecally enriched reactivity, with darker shades corresponding to higher ITS magnitude.

**Supplementary Figure 2: Peptide-level intrathecal antibody synthesis**

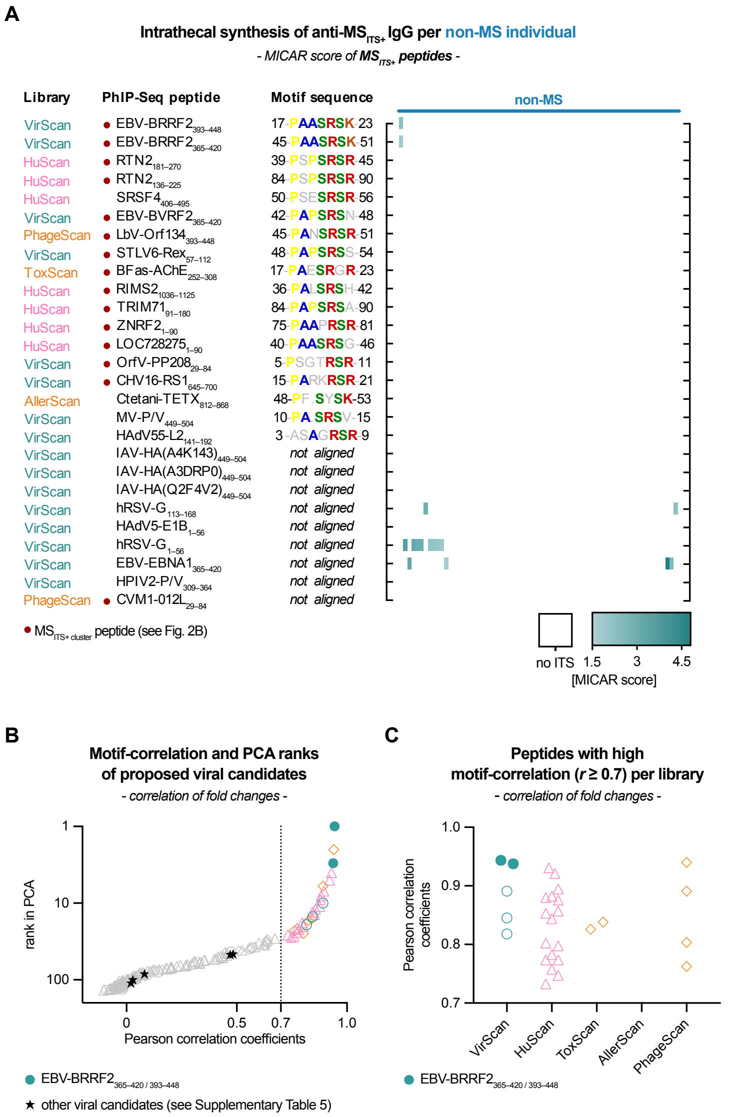


(**A**) Extension of Fig. 2D. Individual-level heatmap of MICAR scores for $MS_{ITS+}$ peptides in individuals with MS (*n* = 40), grouped by $MS_{IC}$ status. White indicates absence of ITS; shades of green indicate magnitude of ITS. Motif sequences represent aligned central peptide regions. Colored residues indicate agreement with the IC-motif; grey residues indicate no agreement.

(**B**) Modified version of Fig. 2E. Correlation analysis of antibody binding among PhIP-Seq library peptides containing a motif with up to one amino acid mismatch to the IC-motif (n = 300 peptides) and binding in at least one sample (n = 137/300 peptides). Each symbol represents one peptide; colors and shapes are as in Fig. 2A. The x-axis shows Pearson correlation coefficients for the five highest-ranking peptides across all samples; the y-axis indicates rank based on the first principal component. Peptides with correlations *r* ≥ 0.7 (*n* = 28) were reanalyzed for sequence alignment. Peptides proposed as potential viral candidates for the IC-motif by previous reports (see Supplementary Table 5) are labeled with a star.

(**C**) PhIP-Seq library origin of motif-correlated peptides ($r \geq 0.7$; $n = 28$), showing that the correlation cluster is predominantly composed of human protein-derived peptides, with EBV BRRF2 as the major viral contributor. Each symbol represents one peptide; colours and shapes are as in Fig. 2A. The x-axis indicates the PhIP-Seq library from which each peptide was derived, and the y-axis shows Pearson correlation coefficients relative to the five highest-ranking peptides across all samples.

**Supplementary Figure 3: Characterization of CSF reactivity across proteomes of motif-correlated MS$_{ITS+\ cluster}$ peptides**

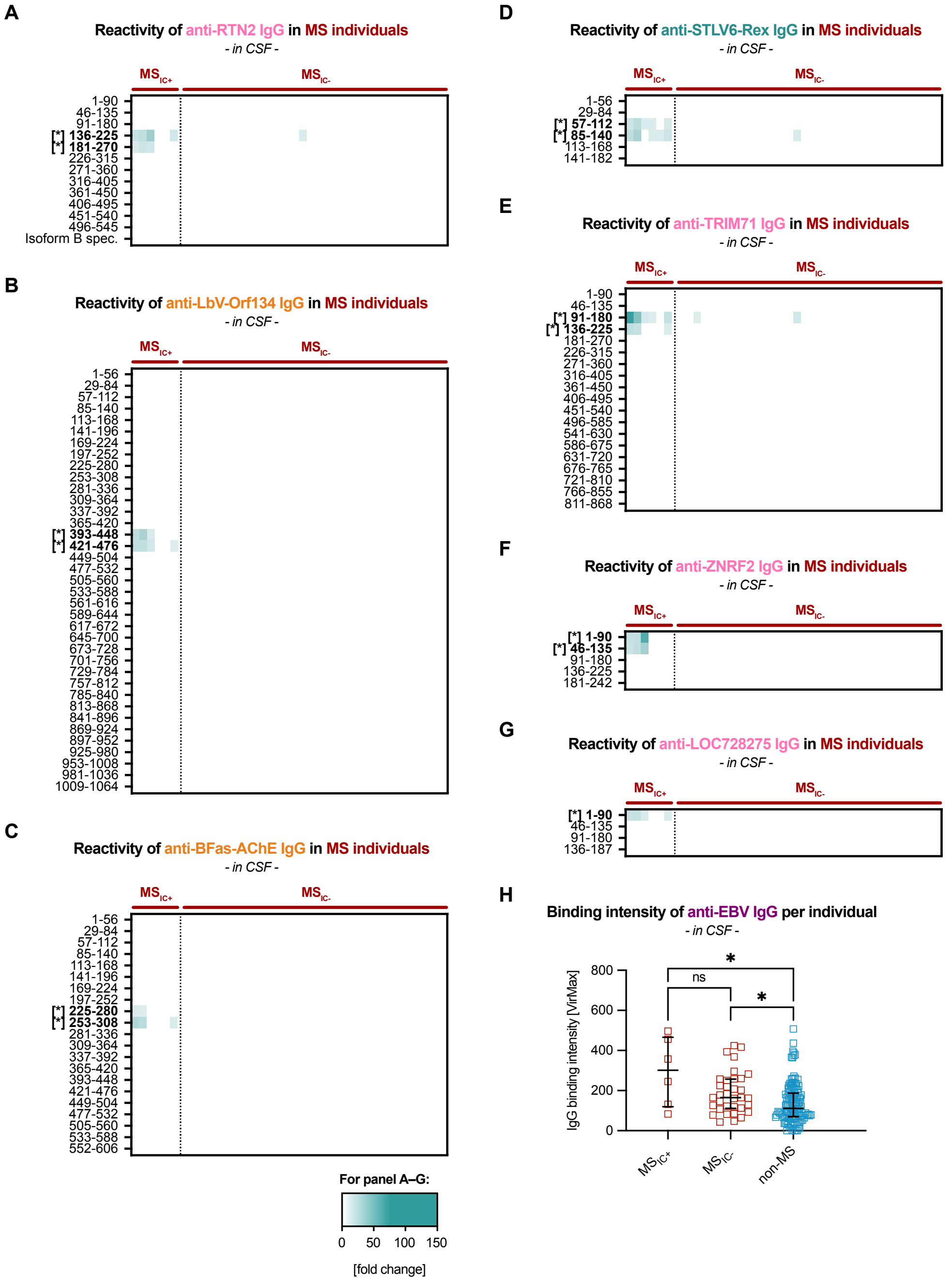


(**A–G**) Individual-level heatmaps of CSF reactivity across proteomes of proteins associated with MS$_{ITS+\ cluster}$ peptides with motif-correlation ($r \geq 0.7$) in addition to Fig. 3C showing CSF reactivity for the EBV BRRF2 protein. Each panel displays one protein, including human proteins RTN2 (A), TRIM71 (E), ZNRF2 (F) and LOC728275 (G) (see Supplementary Table 4 for protein names and associated pathogens). Rows correspond to separate tiles as represented in the PhIP-Seq library. Positions within proteins indicated with an asterisks indicating IC-motif containing peptides. Columns represent individuals stratified by $MS_{IC}$ status ($MS_{IC+}$: *n* = 6, $MS_{IC-}$: *n* = 34). White indicates absence of detectable reactivity; shades of green indicate magnitude of antibody binding intensity.

(**H**) Individual-level EBV-specific CSF antibody binding intensity, measured as VirMax, stratified by $MS_{IC}$ status. Each square represents one individual ($MS_{IC+}$: *n* = 6, $MS_{IC-}$: *n* = 34, non-MS: *n* = 83). Group differences were assessed using a Kruskal–Wallis test (*P* < .01) with Dunn's post-hoc. Horizontal lines indicate the median; whiskers indicate the interquartile range. Illustration is related to Fig. 1F.

#### Supplementary Figure 4: Validation of an ELISA-based serum IgG motif testing protocol

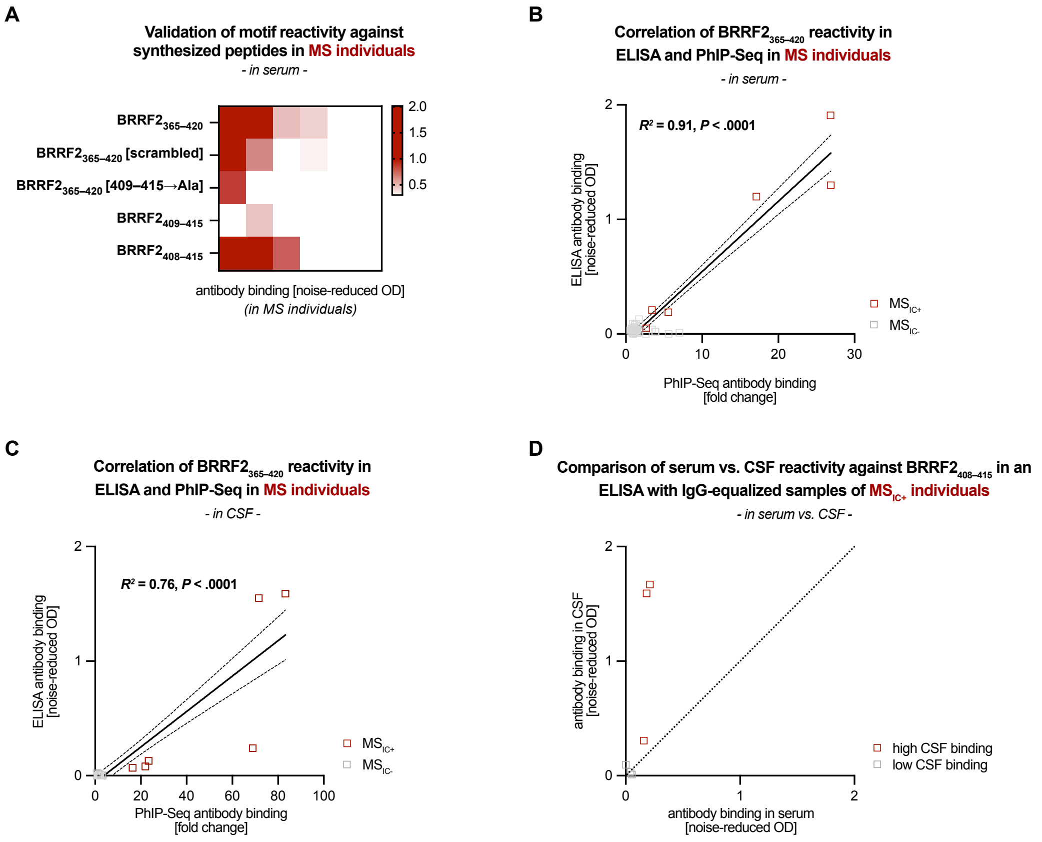


(**A**) Individual-level heatmap of ELISA-based serum IgG reactivity against peptides derived from the 56-mer PhIP-Seq sequence (BRRF2$_{365–420}$), including: the full-length peptide; a variant in which the central PAASRSK motif scrambled to SSPKARA (BRRF2$_{365–420}$[scrambled]); an alanine-substituted variant in which residues 409–415 were replaced by alanines (BRRF2$_{365–420}$[409–415→Ala]; the isolated core motif (BRRF2$_{409–415}$); and an extended variant including one additional N-terminal residue (BRRF2$_{408–415}$). See Supplementary Table 3 for a full list of peptides. Columns represent individuals with MS ($n$ = 6). Shades of Red indicate magnitude of noise-reduced OD.

(**B**) Individual-level ELISA-based serum IgG reactivity against BRRF2$_{365–420}$ correlated with PhIP-Seq binding intensity against the BRRF2$_{365–420}$ tile. Each square represents one individual stratified by MS$_{IC}$ status (MS$_{IC+}$: $n$ = 6, MS$_{IC-}$: $n$ = 34) The x-axis shows PhIP-Seq binding intensity (fold change); the y-axis shows ELISA reactivity (noise-reduced OD). ELISA-based and PhIP-Seq-based serum reactivity to BRRF2$_{365–420}$ are positively associated in a linear regression model ($R^2$ = 0.91, $P$ < .0001). Dashed lines indicate 95% confidence intervals.

(**C**) Corresponding analysis to (B) in CSF. ELISA-based and PhIP-Seq-based CSF reactivity to BRRF2$_{365–420}$ are positively associated in a linear regression model ($R^2$ = 0.76, $P$ < .0001). Dashed lines indicate 95% confidence intervals.

(**D**) Individual-level ELISA-based IgG reactivity to $BRRF2_{408–415}$ in paired, IgG-normalized serum and CSF samples from $MS_{IC+}$ individuals (*n* = 6). Total IgG levels were quantified using a commercial IgG ELISA, and serum samples were diluted to match the corresponding CSF IgG concentration. IgG-matched samples were then assessed for $BRRF2_{408–415}$ reactivity. The x-axis shows noise-reduced OD in serum and the y-axis in CSF. Each square represents one individual; samples with high PhIP-Seq binding intensity to $BRRF2_{365–420}$ in CSF are shown in red (fold change ≥ 25), and those with lower binding intensity in grey (fold change < 25). The dotted line indicates the line of identity (45°); values above the line indicate relatively higher IgG levels of $BRRF2_{408–415}$-specific IgG in CSF compared to serum.

**Supplementary Figure 5: Exploration of EBV-motif co-reactivities, temporal stability and clinical associations**

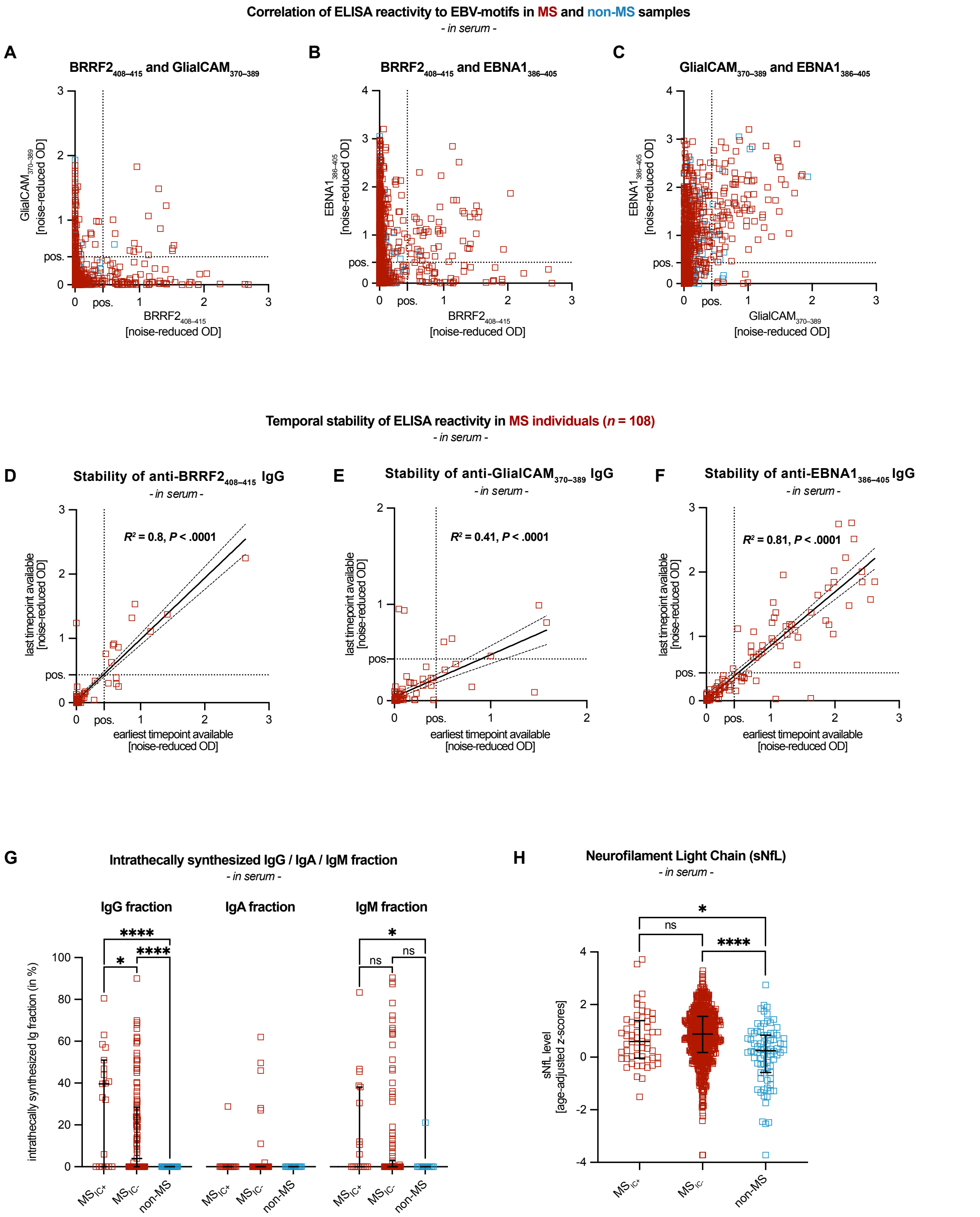


(**A–C**) Individual-level ELISA-based serum IgG reactivity to BRRF2408–415, GlialCAM370–389, and EBNA1386–405. Each square represents one individual (MS: *n* = 909, non-MS: *n* = 311). The x- and y-axes show noise-reduced OD for the indicated peptide pairs.

(**D–F**) Temporal stability of ELISA-based serum IgG reactivity to $BRRF2_{408–415}$, $GlialCAM_{370–389}$, and $EBNA1_{386–405}$ peptides, respectively. Each square represents one MS individual with available longitudinal samples (*n* = 108). The x-axis shows the noise-reduced OD at the earliest available timepoint, and the y-axis shows noise-reduced OD at the last available timepoint. ELISA-based IgG reactivity measurements at both timepoints are positively correlated in a linear regression model for $BRRF2_{408–415}$ ($R^2$ = 0.8, $P$ < .0001), $GlialCAM_{370–389}$ ($R^2$ = 0.41, $P$ < .0001) and $EBNA1_{386–405}$ ($R^2$ = 0.81, $P$ < .0001).

(**G**) Individual-level intrathecal synthesis fractions of IgG, IgA and IgM measurements stratified by $MS_{IC}$ status. Each square represents one individual with varying data availability for IgG ($MS_{IC+}$: *n* = 19, $MS_{IC-}$: *n* = 121, non-MS: *n* = 26), for IgA ($MS_{IC+}$: *n* = 19, $MS_{IC-}$: *n* = 116, non-MS: *n* = 19) and for IgM ($MS_{IC+}$: *n* = 19, $MS_{IC-}$: *n* = 116, non-MS: *n* = 15). Group differences were assessed using a Kruskal–Wallis test (IgG: $P$ < .0001; IgA: $P$ = .55, IgM: P < .05) with Dunn's post-hoc where appropriate. Horizontal lines indicate the median; whiskers indicate the interquartile ranges.

(**H**) Individual-level age-adjusted z-scores for serum Neurofilament Light Chain (sNfL) measurements stratified by $MS_{IC}$ status. Each square represents one individual ($MS_{IC+}$: *n* = 54, $MS_{IC-}$: *n* = 646, non-MS (all HC/OND): *n* = 78). Group differences were assessed using a Kruskal–Wallis test ($P$ < .0001) with Dunn's post-hoc. Horizontal lines indicate the median; whiskers indicate the interquartile ranges.

### SUPPLEMENTARY TABLES

#### Supplementary Table 1: Sample characteristics of the discovery cohort

| Variables | MS | non-MS | non-MS per subgroup[a] | | |
|---|---|---|---|---|---|
| | | | $H^+N^+$ | $H_{only}$ | $N_{only}$ |
| **Number of individuals, n** | 40 | 83 | 13 | 5 | 65 |
| **Sample characteristics** | | | | | |
| Age in years, median (IQR) | 33.5 (28–39) | 32 (22–42) | 64 (38–75) | 32 (24.5–50.5) | 29 (21–35.5) |
| Sex, %Female | 67.5% | 74.7% | 61.5% | 80% | 76.9% |
| Months since symptom onset, median (IQR) | 6 (1–23) | 4 (0–24) | 2 (0–6.5) | 36 (0–184) | 7 (0–24) |
| **Geographic distribution** | | | | | |
| Germany, n (% of total) | 40 (100%) | 81 (97.6%) | 13 (100%) | 3 (60%) | 65 (100%) |
| **CSF parameters** | | | | | |
| CSF WBC in cells/µL, median (IQR) | 10 (5–18) | 3 (1–12) | 5 (2–32.5) | 8 (3–39) | 3 (1–8) |
| CSF Protein in mg/L, median (IQR) | 352 (270–421) | n/a | n/a | n/a | n/a |
| Age-adjusted Qalb, median (IQR) | 0.7 (0.5–0.9) | 0.7 (0.5–1) | 0.8 (0.6–1.6) | 0.4 (0.2–4.2) | 0.7 (0.5–1) |
| CSF-specific Oligoclonal Bands, % | 100% | 48.2% | 69.2% | 40% | 44.6% |
| **Intrathecal Synthesis parameters** | | | | | |
| Intrathecal IgG synthesis in %, median (IQR) | 25 (8.5–46.5) | 0 (0–19.1) | 40.4 (0–45.8) | 0 (0–19.2) | 0 (0–0) |
| Intrathecal IgG synthesis >10%, % | 72.5% | 26.5% | 69.2% | 20% | 18.5% |

[a] As non-MS controls, we used samples from patients with NMDARE or HSE, abbreviated as follows: HSE only ($H_{only}$), NMDARE only ($N_{only}$), HSE and secondary NMDARE ($H^+N^+$)

#### Supplementary Table 2: Sample characteristics of the validation cohort

| | | | non-MS per subgroup | | | |
|---|---|---|---|---|---|---|
| **Variables** | **MS** | **non-MS** | **NMOSD** | **MOGAD** | **HC/OND** | **Data availability per variable[a]** |
| **Number of individuals, n** | 909 | 311 | 78 | 93 | 140 | |
| follow-up measurement available, n | 108 | 0 | 0 | 0 | 0 | |
| **Sample characteristics** | | | | | | |
| Age in years, median (IQR) | 48 (37–57) | 48 (36–59) | 55 (43–63) | 35 (27–48) | 53 (42–60) | [848, 218 (50, 71, 97)] |
| Sex, %Female | 72.9% | 64.1% | 92.2% | 61.1% | 51.5% | [848, 220 (51, 72, 97)] |
| Months since diagnosis, median (IQR) | 108 (48–192) | n/a | 42 (15–111) | 23 (5–64) | n/a | [847, n/a (50, 70, n/a)] |
| **Geographic distribution** | | | | | | |
| Germany, n (% of total) | 149 (16%) | 184 (51%) | 51 (65%) | 73 (78%) | 60 (31%) | [909, 311 (78, 93, 140)] |
| **Clinical characteristics** | | | | | | |
| EDSS, median (IQR) | 3 (1.5–3.5) | 2.75 (1.5–4) | 3.5 (2–4.5) | 2 (1.5–3.5) | n/a | [734, 69 (33, 36, n/a)] |
| Serum NfL in pg/mL, median (IQR) | 11.1 (8.7–15) | 10.8 (7.3–13.9) | n/a | n/a | 10.8 (7.3–13.9) | [700, 79 (n/a, n/a, 79)] |
| Serum NfL age-adjusted z-score, median (IQR) | 0.9 (0.2–1.5) | 0.3 (-0.5–0.9) | n/a | n/a | 0.3 (-0.5–0.9) | [700, 79 (n/a, n/a, 79)] |
| **CSF parameters** | | | | | | |
| CSF WBC in cells/µL, median (IQR) | 5 (3–11) | 7.5 (2.3–13.8) | 3 (1.5–15.5) | 8 (2.5–22) | n/a | [102, 30 (9, 21, n/a)] |
| CSF Protein in mg/L, median (IQR) | n/a | 365 (334–442) | 399 (242–437) | 358 (337–444) | n/a | [n/a, 22 (5, 17, n/a)] |
| Age-adjusted Qalb, median (IQR) | 0.8 (0.6–1) | 0.8 (0.7–1) | 0.8 (0.7–1.2) | 0.8 (0.7–1) | n/a | [99, 17 (3, 14, n/a)] |
| CSF-specific Oligoclonal Bands, % | 89.2% | 16.7% | 17.6% | 13% | n/a | [102, 63 (17, 46, n/a)] |
| **Intrathecal Synthesis parameters** | | | | | | |

| | | | | | | |
|---|---|---|---|---|---|---|
| Intrathecal IgG synthesis in %, median (IQR) | 0 (0–26.4) | 0 (0–0) | 0 (0–0) | 0 (0–0) | n/a | [100, 26 (7, 19, n/a] |
| Intrathecal IgA synthesis in %, median (IQR) | 0 (0–0) | 0 (0–0) | 0 (0–0) | 0 (0–0) | n/a | [95, 19 (3, 16, n/a)] |
| Intrathecal IgM synthesis in %, median (IQR) | 0 (0–0.8) | 0 (0–0) | 0 (0–0) | 0 (0–0) | n/a | [98, 15 (3, 12, n/a)] |

[a] Number of samples with available data for each respective variable. Format: [MS, non-MS (NMOSD, MOGAD, HC/OND)]

Abbreviations: CSF = cerebrospinal fluid; EDSS = Expanded Disability Status Scale; IgA/G/M = immunoglobulin A/G/M; IQR = interquartile range; MS = multiple sclerosis; $MS_{IC}$ = MS immunogenic cluster; NfL = neurofilament light chain; OCB = oligoclonal band; QAlb = albumin quotient; WBC = white blood cell count.

#### Supplementary Table 3: List of synthesized peptides

| Name | Modifications | Length | Sequence |
|---|---|---|---|
| **BRRF2 constructs** | | | |
| $BRRF2_{365–420}$ | His-Tag, Strep-Tag II | 56 + 16 | PPVCPIVSLTASGAKQNRGGMGSLHLAKPEETSPAVSPVCP IASPAASRSKQHCGVHHHHHHSAWSHPQFEK-COOH |
| $BRRF2_{365–420}$ scrambled | His-Tag, Strep-Tag II | 56 + 16 | PPVCPIVSLTASGAKQNRGGMGSLHLAKPEETSPAVSPVCP IASSSPKARAQHCGVHHHHHHSAWSHPQFEK-COOH |
| $BRRF2_{365–420}$ alanine-substituted | His-Tag, Strep-Tag II | 56 + 16 | PPVCPIVSLTASGAKQNRGGMGSLHLAKPEETSPAVSPVCP IASAAAAAAAQHCGVHHHHHHSAWSHPQFEK-COOH |
| $BRRF2_{409–415}$ | His-Tag, Strep-Tag II | 7 + 16 | PAASRSKHHHHHHSAWSHPQFEK-COOH |
| $BRRF2_{408–415}$ | His-Tag, Strep-Tag II | 8 + 16 | SPAASRSKHHHHHHSAWSHPQFEK-COOH |
| **EBNA1-mimic constructs** | | | |
| $GlialCAM_{370–389}$ | His-Tag, Strep-Tag II, pSer376 | 20 + 16 | ATGRTHSpSPPRAPSSPGRSRHHHHHHSAWSHPQFEK-COOH |
| $EBNA1_{386–405}$ | His-Tag, Strep-Tag II | 20 + 16 | SQSSSSGSPPRRPPPGRRPFHHHHHHSAWSHPQFEK-COOH |
| **Human protein constructs** | | | |
| $TRIM71_{173–180}$ | His-Tag, Strep-Tag II | 8 + 16 | PPAPSRSAHHHHHHSAWSHPQFEK-COOH |
| $RTN2_{218–225}$ | His-Tag, Strep-Tag II | 8 + 16 | TPSPSRSRHHHHHHSAWSHPQFEK-COOH |
| $ZNRF2_{74–81}$ | His-Tag, Strep-Tag II | 8 + 16 | APAAPRSRHHHHHHSAWSHPQFEK-COOH |
| $RIMS2_{1080–1087}$ | His-Tag, Strep-Tag II | 8 + 16 | SPALSRSHHHHHHHSAWSHPQFEK-COOH |
| **Technical constructs** | | | |
| His-Strep-Tag II | | 16 | HHHHHHSAWSHPQFEK-COOH |

**Supplementary Table 4: List of $MS_{ITS+}$ peptides**

| | | | | | Network Analysis | |
|---|---|---|---|---|---|---|
| **Peptide Name** | **Library** | **Species** | **UniProt ID** | **Sequence** | **Node Centrality** | **Cluster** |
| EBV-BRRF2$_{393–448}$ | VirScan | Epstein-Barr virus | Q1HVF8 | PEETSPAVSPVCPIASPAASRSKQHCGVTGSSQAA PSSSSVAPVASLSGDLEEEEE | 6.00 | 1 |
| EBV-BRRF2$_{365–420}$ | VirScan | Epstein-Barr virus | Q1HVF8 | PPVCPIVSLTASGAKQNRGGMGSLHLAKPEETSPA VSPVCPIASPAASRSKQHCGV | 4.05 | 1 |
| RTN2$_{181–270}$ | HuScan | Homo sapiens | O75298 | TGEAGEELDLRLRLAQPSSPEVLTPQLSPGSGTPQ AGTPSPSRSRDSNSGPEEPLLEEEEKQWGPLEREP VRGQCLDSTDQLEFTVEPRL | 5.82 | 1 |
| RTN2$_{136–225}$ | HuScan | Homo sapiens | O75298 | PLEDLRLRLDHLGWVARGTGSGEDSSTSSSTPLED EEPQEPNRLETGEAGEELDLRLRLAQPSSPEVLTP QLSPGSGTPQAGTPSPSRSR | 7.05 | 1 |
| SRSF4$_{406–495}$ | HuScan | Homo sapiens | Q08170 | SPSRSVSKEREHAKSESSQREGRGESENAGTNQE TRSRSRSNSKSKPNLPSESRSRSKSASKTRSRSKS RSRSASRSPSRSRSRSHSRS | 0.00 | 4 |
| EBV-BVRF2$_{365–420}$ | VirScan | Epstein-Barr virus | Q1HVC7 | DAHTYHPHPHPPPAYFGLPGLFGPPPPVPPYYGSH LRADYVPAPSRSNKRKRDPEE | 6.16 | 1 |
| LbV-Orf134$_{393–448}$ | PhageScan | Phage (Lactobacillus virus LP65) | YP_164769.1 | KKVNPKGSWSKSRKTSENGEPDWTPYILAIIDTEHK LNSSSQSDPANSRSRGSASS | 4.69 | 1 |
| STLV6-Rex$_{57–112}$ | VirScan | Simian T-cell lymphotropic virus | D1MNA2 | PPGYIGTPYWPPVLNTRSPGTPSMDALSARLYNTLS LASPPSPPKELPAPSRSSPR | 5.86 | 1 |
| BFas-AChE$_{252–308}$ | ToxScan | Bungarus fasciatus | Q92035 | AILQSGGPNAPWATVTPAESRGRAALLGKQLGCHF NNDSELVSCLRSKNPQELIDE | 4.87 | 1 |
| RIMS2$_{1036–1125}$ | HuScan | Homo sapiens | Q9UQ26 | LLERTTTRSRSTERPDTNLMRSMPSLMTGRSAPPS PALSRSHPRTGSVQTSPSSTPVAGRRGRQLPQLPP KGTLDRKAGGKKLRSTVQRS | 6.47 | 1 |
| TRIM71$_{91–180}$ | HuScan | Homo sapiens | Q2Q1W2 | CPVCDQKVVLAEAAGMDALPSSAFLLSNLLDAVVAT ADEPPPKNGRAGAPAGAGGHSNHRHHAHHAHPRA SASAPPLPQAPQPPAPSRSA | 7.67 | 1 |
| ZNRF2$_{1–90}$ | HuScan | Homo sapiens | Q8NHG8 | MGAKQSGPAAANGRTRAYSGSDLPSSSSGGANGT AGGGGGARAAAAGRFPAQVPSAHQPSASGGAAAA AAAPAAPAAPRSRSLGGAVGSV | 6.33 | 1 |
| LOC728275$_{1–90}$ | HuScan | Homo sapiens | n/a | MPAHHDPQILDGDSGEEAETTTPQCGTEPPSSGLKI SAIPAASRSGPGASPTVNTDPGLLPKWGREQAASR ETGWKGLSPTTASLPLLRG | 8.56 | 1 |
| OrfV-PP208$_{29–84}$ | VirScan | Parapoxvirus / Orf virus | F1AX61 | TSRTPSGTRSRAKYRAGTGSGARPCGPWASACGP CSCWSWRSLRAMTASRPWPALW | 2.87 | 1 |

| | | | | | | |
|---|---|---|---|---|---|---|
| CHV16-RS1$_{645–700}$ | VirScan | Cercopithecine herpesvirus 16 | Q2QBA3 | SSSPPRAPAPAARPPARKRSRPARPRPPGAPADDD DDNGGGRARAPGPRPRPAALT | 6.75 | 1 |
| Ctetani-TETX$_{812–868}$ | ToxScan | Clostridium tetani | P04958 | AKKQLLEFDTQSKNILMQYIKANSKFIGITELKKLESK INKVFSTPIPFSYSKNLD | 0.00 | 9 |
| MV-P/V$_{449–504}$ | VirScan | Measles virus | P35974 | AVGFVPDTGPASRSVIRSIIKSSRLEEDRKRYLMTLL DDIKGANDLAKFHQMLMKI | 0.00 | 8 |
| HAdV55-L2$_{141–192}$ | VirScan | Human adenovirus type 55 | C7SRT6 | SGASAGRSRRQAAAVAAATIADMAQTRRGNVYWV RDAATGQRVPVRTRPPRT | 0.00 | 5 |
| IAV-HA(A4K143)$_{449–504}$ | VirScan | Influenza A virus | A4K143 | ERTLDFHDSNVKNLYEKVKNQLRNNAKEIGNGCFE FYHKCDNECMESVKNGTYDYP | 1.93 | 2 |
| IAV-HA(A3DRP0)$_{449–504}$ | VirScan | Influenza A virus | A3DRP0 | ERTLDFHDSNVKNLYEKVKNQLKNNAKEIGNGCFEF YHKCNNECMESVKNGTYDYP | 1.91 | 2 |
| IAV-HA(Q2F4V2$_{449–504}$ | VirScan | Influenza A virus | Q2F4V2 | NERTLDFHDSNVKNLYDKVRLQLRDNAKELGNGCF EFYHKCDNECMESVRNGTYDY | 1.90 | 2 |
| hRSV-G$_{113–168}$ | VirScan | Human respiratory syncytial virus | E9NW54 | PCSICSNNPTCWAICKRIPNKKPGKKTTTKPTKKPTI KTTKKDPKPQTTKPKEVLT | 0.00 | 10 |
| HAdV5-E1B$_{1–56}$ | VirScan | Human adenovirus C serotype 5 | P03243 | MERRNPSERGVPAGFSGHASVESGCETQESPATV VFRPPGDNTDGGAAAAAGGSQA | 0.00 | 6 |
| hRSV-G$_{1–56}$ | VirScan | Human respiratory syncytial virus | C1K8X5 | PAQTNRPSTKPRPKNPPKKPKDDYHFEVFNFVPCSI CGNNQLCKSICKTIPSNKPK | 0.00 | 7 |
| EBV-EBNA1$_{365–420}$ | VirScan | Epstein-Barr virus | Q1HVF7 | SRERARGRGRGRGEKRPRSPSSQSSSSGSPPRRP PPGRRPFFHPVAEADYFEYHQE | 0.53 | 3 |
| HPIV2-P/V$_{309–364}$ | VirScan | Human parainfluenza type 2 virus | P23055 | SGGFTAEGSDMISMDELARPTLSSTKRITRKPESKK DLTGIKLTLMQLANDCISRP | 0.53 | 3 |
| CVM1-012L$_{29–84}$ | PhageScan | Phage (Paramecium bursaria Chlorella virus) | AGE51696 | TTSDVKMPSKFTVDKWKYNKSVLPFYDTRDYEESR QGLMATPAYKQIKDADGKVVW | 7.16 | 1 |

#### Supplementary Table 5: Motif score correlation

| | | | | | IC-motif correlation analysis | |
|---|---|---|---|---|---|---|
| **Peptide Name** | **Library** | **Associated Species** | **UniProt ID** | **IC-motif Pattern** | **Correlation Coefficient** | **FPC Rank[a]** |
| **Peptides with IC-motif correlation scores ≥ 0.7** | | | | | | |
| EBV-BRRF2$_{393–448}$ | VirScan | Epstein-Barr virus | Q1HVF8 | PAASRSK | 0.943608 | 1 |
| LbV-Orf134$_{421–477}$ | HuScan | Homo sapiens | YP_164769.1 | PANSRSR | 0.9398 | 2 |
| EBV-BRRF2$_{365–420}$ | VirScan | Epstein-Barr virus | Q1HVF8 | PAASRSK | 0.937504 | 3 |
| KRT75$_{1–91}$ | HuScan | Homo sapiens | O95678 | PAAGRSR | 0.930814 | 4 |
| LOC728275$_{1–91}$ | HuScan | Homo sapiens | n/a | PAASRSG | 0.921057 | 5 |
| CFAP97$_{46–136}$ | HuScan | Homo sapiens | Q9P2B7 | PASSRSK | 0.894666 | 7 |
| LbV-Orf134$_{393–449}$ | PhageScan | Phage (Lactobacillus virus LP65) | YP_164769.1 | PANSRSR | 0.890959 | 6 |
| STLV6-Rex$_{57–112}$ | VirScan | Simian T-cell lymphotropic virus | D1MNA2 | PAPSRSS | 0.89081 | 10 |
| RTN2$_{181–271}$ | HuScan | Homo sapiens | O75298 | PSPSRSR | 0.882711 | 8 |
| LOC100287739$_{1–91}$ | HuScan | Homo sapiens | n/a | PSRSRSR | 0.880118 | 9 |
| K2C75$_{1–56}$ | HuScan | Homo sapiens | O95678 | PAAGRSR | 0.87583 | 11 |
| TRIM71$_{136–226}$ | HuScan | Homo sapiens | Q2Q1W2 | PAPSRSA | 0.857275 | 12 |
| TRIM71$_{91–181}$ | HuScan | Homo sapiens | Q2Q1W2 | PAPSRSA | 0.85335 | 14 |
| STLV6-Rex$_{85–140}$ | VirScan | Simian T-cell lymphotropic virus | D1MNA2 | PAPSRSS | 0.845473 | 15 |
| CFAP97$_{91–181}$ | HuScan | Homo sapiens | Q9P2B7 | PASSRSK | 0.844051 | 13 |
| BFas-AChE$_{252–308}$ | ToxScan | Bungarus fasciatus | Q92035 | PAESRGR | 0.838344 | 16 |
| CAlb-WOR2_CANAL$_{56–112}$ | ToxScan | Candida albicans | Q5ANB1 | PSPGRSK | 0.82613 | 17 |

| | | | | | | |
|---|---|---|---|---|---|---|
| HHV1$_{29–81}$ | VirScan | Human herpesvirus 1 | Q6VB60 | PAASRSV | 0.817885 | 19 |
| StrmV-VWB$_{57–113}$ | PhageScan | Phage (Streptomyces phage VWB) | NP_958294.1 | PATSRSK | 0.803416 | 25 |
| RTN2$_{136–226}$ | HuScan | Homo sapiens | O75298 | PSPSRSR | 0.802219 | 20 |
| ZNRF2$_{46–136}$ | HuScan | Homo sapiens | Q8NHG8 | PAAPRSR | 0.797741 | 18 |
| FOXO6$_{406–496}$ | HuScan | Homo sapiens | n/a | PAPSRSA | 0.782512 | 22 |
| DENND4C$_{1036–1126}$ | HuScan | Homo sapiens | Q5VZ89 | PAVSRSK | 0.773939 | 24 |
| FOXO6$_{319–388}$ | HuScan | Homo sapiens | n/a | PAPSRSA | 0.773893 | 21 |
| EMas$_{29–85}$ | PhageScan | Eisenbergiella massiliensis | WP_117531915.1 | PAPPRSK | 0.762369 | 23 |
| TYSND1$_{46–136}$ | HuScan | Homo sapiens | Q2T9J0 | PAPSRGR | 0.757875 | 27 |
| FOXO6$_{451–541}$ | HuScan | Homo sapiens | n/a | PAPSRSA | 0.747287 | 28 |
| CLASRP$_{496–586}$ | HuScan | Homo sapiens | Q8N2M8 | LSPSRSR, PSQSRSR, PSPSQSR, PSRSRSL | 0.732510 | 26 |
| **Other viral candidates hypothesized in prior work**[b] | | | | | | |
| EBV-gM$_{309–364}$ | VirScan | Epstein-Barr virus | P03215 | PSPGRNR | 0.483267 | 46 |
| EBV-gM$_{337–392}$ | VirScan | Epstein-Barr virus | P03215 | PSPGRNR | 0.470714 | 47 |
| REBOV-VP30$_{1–56}$ | VirScan | Reston ebolavirus (Phillipines-96) | Q91DD6 | PSRSRSL | 0.081692 | 84 |
| HCV6h$_{1093–1148}$ | VirScan | Hepatitis C virus genotype 6h | O92532 | PAGARSL | 0.028485 | 100 |
| REBOV-VP30$_{1–56}$ | VirScan | Reston ebolavirus (Reston-89) | Q8JPX6 | PSRSRSL | 0.019791 | 110 |

[a] Rank in the first principal component of a PCA on relative binding scores of 137/300 peptides which bound in at least one sample across all CSF and serum samples from MS and non-MS cohorts. See *Materials and methods* and Fig. 2A.
[b] see Zamecnik CR, Sowa GM, Abdelhak A, et al. An autoantibody signature predictive for multiple sclerosis. *Nat Med*. 2024;30(5):1300-1308. doi:10.1038/s41591-024-02938-3

**Supplementary Table 6: Association of ELISA-based anti-motif IgG seropositivity with MS diagnosis**

| Predictor | Odds ratio | 95% Confidence Interval | *P*-value | Samples included[a] | Number of events[a] |
|---|---|---|---|---|---|
| **Unadjusted logistic regression** | | | | | |
| $BRRF2_{408–415}$ | 14.1 | 4.4–86.04 | .0002 | 1220 | 909 |
| $GlialCAM_{370–389}$ | 2.81 | 1.58–5.46 | .001 | 1220 | 909 |
| $EBNA1_{386–405}$ | 2.23 | 1.69–2.95 | < .0001 | 1220 | 909 |
| **Unadjusted Firth penalized logistic regression** | | | | | |
| $BRRF2_{408–415}$ | 11.4 | 3.91–54.9 | < .0001 | 1220 | 909 |
| $GlialCAM_{370–389}$ | 2.71 | 1.54–5.21 | .0003 | 1220 | 909 |
| $EBNA1_{386–405}$ | 2.22 | 1.68–2.94 | < .0001 | 1220 | 909 |
| **Adjusted Firth penalized logistic regression**[b] | | | | | |
| $BRRF2_{408–415}$ | 9.48 | 3.12–46.8 | < .0001 | 1064 | 846 |
| $GlialCAM_{370–389}$ | 3.61 | 1.7–8.63 | .0005 | 1064 | 846 |
| $EBNA1_{386–405}$ | 2.54 | 1.76–3.72 | < .0001 | 1064 | 846 |

[a] Adjusted analyses were restricted to complete cases across all included variables, thus number of included samples and events (samples from individuals with MS diagnosis) are reduced in comparison to Fig. 4A.

[b] Adjusted for age, sex and country of sampling.

**Supplementary Table 7: Associations between ELISA-based anti-motif IgG seropositivity measures**

| Predictor | Odds ratio | 95% Confidence Interval | *P*-value | Samples included[a] | Number of events[a] |
|---|---|---|---|---|---|
| **Adjusted Firth penalized logistic regression on EBNA1$_{386–405}$[b]** | | | | | |
| BRRF2$_{408–415}$ | 1.45 | 0.87–2.44 | .1568 | 1064 | 442 |
| GlialCAM$_{370–389}$ | 19.72 | 9.86–45.94 | < .0001 | 1064 | 442 |
| **Adjusted Firth penalized logistic regression on BRRF2$_{408–415}$[b]** | | | | | |
| GlialCAM$_{370–389}$ | 1.81 | 0.9–3.46 | .0929 | 1064 | 74 |
| EBNA1$_{386–405}$ | 1.45 | 0.86–2.43 | .162 | 1064 | 74 |

[a] Adjusted analyses were restricted to complete cases across all included variables, thus number of included samples and events (seropositive samples) are reduced in comparison to Fig. 4A.

[b] Adjusted for age, sex, country of sampling and disease group (MS / non-MS)

#### Supplementary Table 8: Human Protein List[a]

| Gene name | Gene Product | UniProt ID | Predicted Location | Tissue profile | Tissue specificity | Single cell type specificity | Brain specificity | Brain expression cluster | Phenotype from variants (MIM #) |
|---|---|---|---|---|---|---|---|---|---|
| RTN2 | Reticulon-2 / Neuroendocrine-specific protein-like | O75298 | Membrane | Cytoplasmic expression in muscle cells | Skeletal muscle, tongue | Extravillous trophoblasts, Cytotrophoblasts, Platelets, Syncytiotrophoblasts, Migrating cytotrophoblasts, Myonuclei | Low | Neurons & Synapses - Synaptic function (mainly) | Spastic paraplegia 12 (604805) |
| KRT75 (K2C75) | Keratin, type II cytoskeletal 75 | O95678 | Intracellular | Distinct cytoplasmic expression in the outer root sheath of hair follicles | Skin | Pancreatic endocrine cells, Suprabasal keratinocytes, Late spermatids, Basal keratinocytes, Serous glandular cells, Basal respiratory cells | n/a | n/a | Loose anagen hair syndrome (600628), pseudofolliculitis barbae (612318) |
| ZNRF2 | E3 ubiquitin-protein ligase ZNRF2 | Q8NHG8 | Intracellular | General cytoplasmic expression | Low | Microglial cells | Low | Macrophages & Microglia - Immune response (mainly) | n/a |
| CFAP97 | Cilia- and flagella-associated protein 97 | Q9P2B7 | Intracellular | Ubiquitous cytoplasmic and membranous expression | Low | Early spermatids | Low | Non-specific - Transcription (mainly) | n/a |
| TRIM71 | E3 ubiquitin-protein ligase TRIM71 | Q2Q1W2 | Intracellular | n/a | Testis | Extravillous trophoblasts, Spermatogonia, Horizontal cells, Alveolar cells type 2, Oocytes, Alveolar cells type 1 | Low | Neurons - Mixed function (mainly) | Hydrocephalus, congenital 4; HYC4 (618667) |
| DENND4C | DENN domain-containing protein 4C | Q5VZ89 | Membrane, Intracellular (different isoforms) | Cytoplasmic expression in most tissues | Low | Low | Low human brain regional specificity | Non-specific - Transcription (mainly) | n/a |

| | | | | | | | | | |
|---|---|---|---|---|---|---|---|---|---|
| FOXO6 | Forkhead box protein O6 | n/a | Intracellular | n/a | n/a | Granulosa cells, Mesothelial cells, Skeletal myocytes | Low | Choroid plexus - Cilium (mainly) | n/a |
| TYSND1 | Peroxisomal leader peptide-processing protease | Q2T9J0 | Intracellular | Cytoplasmic expression in most tissues | Low | Late spermatids, Early spermatids, Spermatocytes | Low | Non-specific - Transcription (mainly) | n/a |
| CLASRP | CLK4 associating serine/arginine rich protein | Q8N2M8 | Intracellular | Nuclear expression in several tissues | Low | Low | Low | Non-specific – Immune response (mainly) | n/a |

[a] For human motif-containing proteins with correlation scores to IC-motif $r \geq 0.7$ for the respective motif-containing peptide(s), data on protein and RNA expression were extracted from the Human Protein Atlas (proteinatlas.org). Data on protein-related human diseases derives from OMIM database (omim.org). Hypothetical proteins LOC100287739 and LOC728275 were excluded from extraction due to the absence of functional characterization.
